# Evidence of 14-3-3 proteins contributing to kinetochore integrity and chromosome congression during mitosis

**DOI:** 10.1101/2023.12.24.573259

**Authors:** Guhan Kaliyaperumal Anbalagan, Prakhar Agarwal, Santanu Kumar Ghosh

## Abstract

The 14-3-3 family of proteins are conserved across eukaryotes and serve myriad important regulatory functions of the cell. Homo/heterodimers of these protein homologs, majorly recognize their ligands via conserved motifs from a plethora of cellular proteins to modulate the localization and functions of those effector ligands. In most of the genetic backgrounds of *Saccharomyces cerevisiae*, disruption of both 14-3-3 homologs (Bmh1 and Bmh2) are either lethal or survive with severe growth defects showing gross chromosomal missegregation and prolonged cell cycle arrest. To elucidate their contributions to chromosome segregation, in this work we investigated their centromere/kinetochore-related functions. Analysis of appropriate deletion mutants shows that Bmh isoforms have cumulative and unshared isoform-specific contributions in maintaining the proper integrity of the kinetochore ensemble. Consequently, *bmh* mutant cells exhibited perturbations in kinetochore-microtubule (KT-MT) dynamics, characterized by kinetochore declustering, mis-localization of kinetochore proteins, and Mad2-mediated transient G2/M arrest. These defects also caused an asynchronous chromosome congression in *bmh* mutants during metaphase. In summary, this report advances the knowledge on contributions of budding yeast 14-3-3 proteins in chromosome segregation by demonstrating their roles in kinetochore integrity and chromosome congression.

## Introduction

The 14-3-3 are a family of highly conserved low molecular weight acidic proteins (Aitken et al., 1992; Martens et al., 1992), known for their myriad functions in eukaryotic cells (Kumar, 2017). Though there is no reported catalytic activity, these proteins generally function by recognizing a highly conserved motif (containing phosphorylated serine/threonine) present on a plethora of proteins to alter their activity, stability, sub-cellular localization, and ability to physically interact with other proteins (Braselmann and McCormick, 1995; Muslin et al., 1996; Yaffe et al., 1997; Lopez-Girona et al., 1999; Sluchanko and Gusev, 2017; Gavade et al., 2022). Other than phosphorylated proteins, 14-3-3 proteins also displayed a strong affinity with non-phosphorylated proteins based on a conserved amphipathic groove (Masters et al., 1999). Among eukaryotes, the number of structurally conserved 14-3-3 isoforms varies with species; the ectopic expression of such isoforms can either partially or completely provide inter-species functional complementation (Van Heusden et al., 1995; Kumar, 2017). Despite such functional complementation among 14-3-3 isoforms, there are unshared functions of individual isoforms as reported elsewhere (Wilkert et al., 2005; Herod et al., 2022).

In budding yeast, there are two 14-3-3 isoforms, namely Bmh1 and Bmh2, that were characterized as having high sequence homology with a mammalian isoform, epsilon (van Heusden et al., 1992; Gelperin et al., 1995; Van Heusden et al., 1995). Bmh1 and Bmh2 exist as homo and heterodimers, of which heterodimers are more prevalent within the cell (Chaudhri et al., 2003). Deletion mutants of either isoform show no impact on cell growth under standard growth conditions, but the double deletion is lethal in most genetic backgrounds of budding yeast (van Heusden et al., 1992); however, in certain genetic backgrounds such as SK1 and ∑1278, the double mutants are viable with severe growth defects due to compromised metabolism of certain carbon sources (Roberts et al., 1997; Slubowski et al., 2014; Kumar, 2018). The viable genetic background strains might have evolved to have an alternative pathway to alleviate the defects caused by the *bmh* double deletions. For instance, over-expression of Ras/cAMP-dependent protein kinase (Tpk1) partially rescued the cell death caused by *bmh* double deletions (Gelperin et al., 1995). Perhaps this is the reason why Bmh proteins are dispensable for cell viability in ∑1278 strain background where Ras/cAMP signaling is hyperactive (Stanhill et al., 1999). Nevertheless, comparing the functions of Bmh proteins from different genetic background strains may help in understanding their unshared and cumulative contributions in budding yeast; therefore, we assessed cumulative functions of Bmh isoforms in SK1 genetic background strains where *bmh* double mutant is viable and unshared functions in W303 background strains where *bmh* double mutant is lethal.

Consistent with their diverse functions across the cell cycle, the Bmh proteins are localized to almost all the sub-cellular locations throughout the mitotic and meiotic cell cycle stages (Slubowski et al., 2014; Kumar, 2018). Their functions during S phase in DNA replication and damage repair have been demonstrated by several studies (Lottersberger et al., 2003; Usui and Petrini, 2007; Yahyaoui et al., 2007; Yahyaoui and Zannis-Hadjopoulos, 2009). The M phase functions argue for their roles in the process of chromosome segregation. For instance, Bmh proteins are involved in spindle-assembly and –position checkpoint mediated cell cycle arrest and hence the mutants are sensitive to spindle damage and spindle misalignment, respectively (Grandin and Charbonneau, 2008; Caydasi et al., 2014). Earlier, we observed a change in the overall cellular protein concentration ratio of Bmh1 and Bmh2 between mitotic and meiotic cells (Kumar et al., 2014; Kumar, 2018), indicating that these proteins may also influence certain events of chromosome segregation that takes place differently in mitosis and meiosis. Notably, there are several lines of evidence depicting that Bmh isoforms have unshared functions in chromosome segregation. For instance, the function of Bmh1 but not Bmh2, has been shown in silencing of spindle-assembly checkpoint (SAC) by facilitating centromeric localization of an intermediate filament protein, Fin1 (Bokros et al., 2016, 2021). Bmh1 acts as a component of spindle position checkpoint (SPOC) that monitors proper orientation of the spindle along the mother-bud axis, ensuring successful disjunction of duplicated SPBs (Caydasi et al., 2014). Recently, Bmh1 has been shown to contribute to chromosome compaction through centromeric recruitment of a histone deacetylase, Hst2 (Jain et al., 2021).

Given Bmh proteins’ role in chromosome segregation through microtubule spindle and chromosome morphogenesis, we hypothesize that these proteins may also influence kinetochore function due to following observations. First, the anti-microtubule drug sensitivity of the *bmh* mutants (Grandin and Charbonneau, 2008), is also shared by the kinetochore protein mutants (Sanyal et al., 1998; Poddar et al., 1999; Ghosh et al., 2001). Second, centromere localized Aurora B kinase (Ipl1) phosphorylates histone H3 at serine 10 that is recognized by Bmh1, which in turn recruits Hst2 at the centromeres to remove H4K16 acetylation required to initiate chromosome compaction from the centromeres in *cis* (Jain et al., 2021). Third, the kinetochore protein Fin1 (Akiyoshi et al., 2009) was shown to have physical interaction with Bmh1 and Bmh2 proteins *in-vitro* (Bokros et al., 2016). Forth, *in-silico* analyses revealed that several kinetochore proteins harbour Bmh1/2 interaction motifs and finally, in a high-throughput study the kinetochore protein Iml3 has been co-purified with Bmh1(Kakiuchi et al., 2007).

To examine the hypothesis, we investigated the functions of the Bmh proteins in kinetochore function and observed that they promote high-fidelity chromosome segregation perhaps by contributing to the maintenance of kinetochore ensemble facilitating a stable KT-MT dynamics. In addition, we provide evidence of Bmh proteins’ role in synchronous chromosome congression during metaphase. Thus, this work reports yet additional functions of Bmh proteins related to faithful chromosome segregation.

## Results

### Bmh proteins contribute to faithful chromosome segregation

Budding yeast cells with mutations in proteins involved in KT-MT related functions of chromosome segregation show hypersensitivity to sub-lethal concentrations of microtubule-destabilizing drugs (Guarente, 1993). To elucidate the contribution of Bmh proteins in similar functions in the context of chromosome segregation, we compared the drug (benomyl) sensitivity of the *bmh* single and double mutants to wild-type and the non-essential kinetochore mutants (*ctf19Δ* and *iml3Δ*) (Fig 1A). Serial dilutions of all the mid-log grown strains were spotted on no-drug (DMSO) and benomyl-containing YEPD plates. As the mitotic growth rate of *bmh1Δ bmh2Δ* double mutant strain was slow as observed in earlier studies (Kumar, 2018), a 2-fold cell concentration of *bmh1Δ bmh2Δ* double mutant was used for normalization purposes in the spotting assays. With increasing concentrations of benomyl, *bmh2Δ* mutant cells showed a growth defect similar to the previously reported *iml3Δ* mutant (Ghosh et al., 2001), while the *bmh1Δ* mutant showed hardly any defect (Fig 1A). The benomyl sensitivity of the *bmh2Δ* mutant is not specific to the strain background, as similar results were obtained in other genetic backgrounds (Fig S1). Noticeably, the severity of benomyl sensitivity varied among different genetic backgrounds of budding yeast (compare Fig 1A with S1). The hypersensitivity of *bmh* single mutants to microtubule destabilizing drugs was observed elsewhere (Grandin and Charbonneau, 2008). However, the *bmh1Δ bmh2Δ* double mutant showed a synergistic growth defect compared to the *bmh* single mutants (Fig 1A) indicating possible unshared functions of Bmh1 and Bmh2 with respect to chromosome segregation. In coherence with our observation, non-redundant functions of 14-3-3 isoforms were reported earlier in higher eukaryotes (Wilkert et al., 2005).

**Figure 1:**
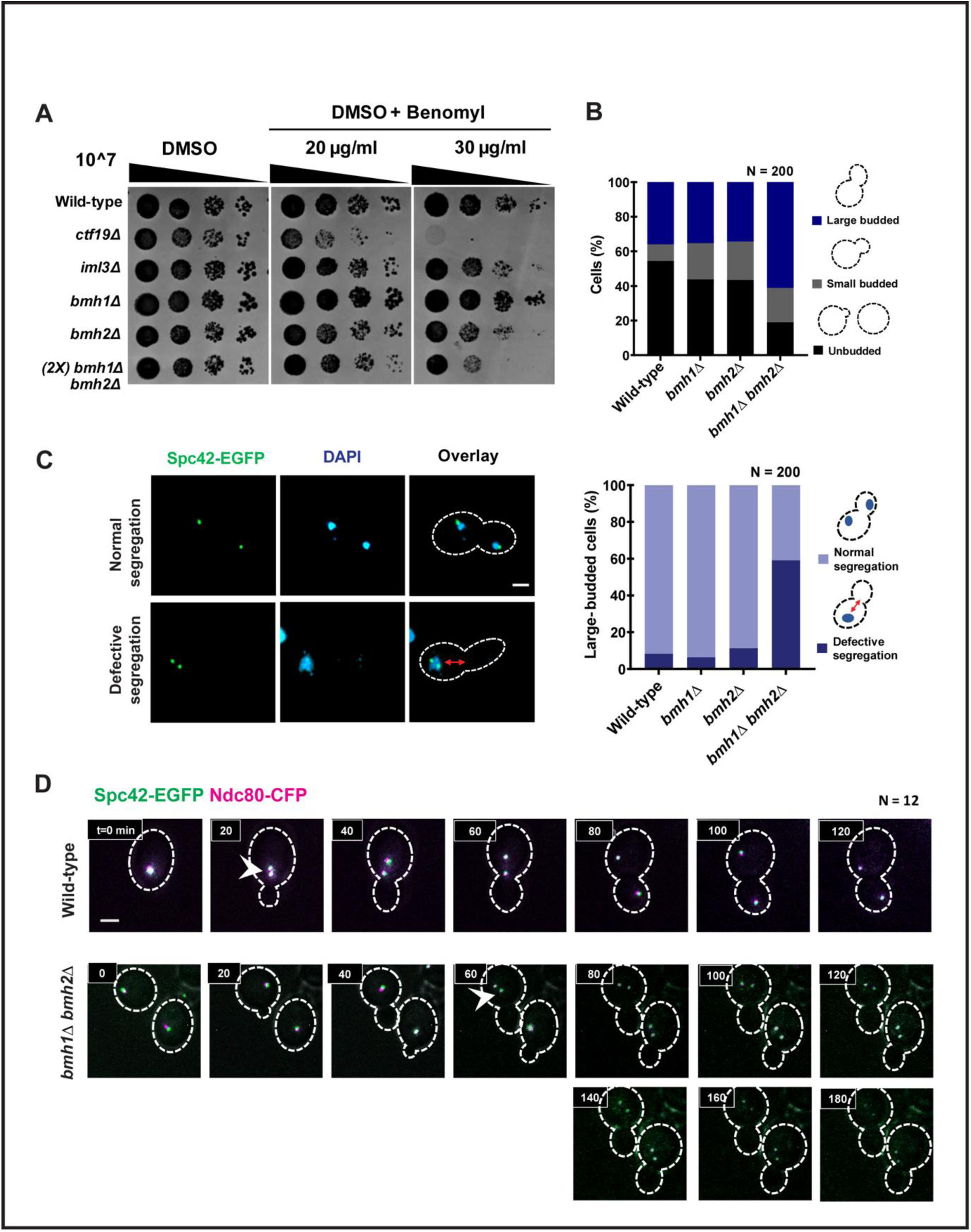
Defective chromosome segregation in the absence of Bmh proteins. (**A**) Hyper-sensitivity of the *bmh* mutants to the microtubule-disrupting drug. Wild-type, *bmh* single (*bmh1Δ, bmh2Δ),* and double (*bmh1Δ bmh2Δ*) mutant cells were spotted on control (DMSO) and indicated concentrations of drug (DMSO+Benomyl) containing YEPD plates. The non-essential kinetochore mutants (*ctf19Δ* and *iml3Δ*) were taken as positive controls for benomyl hyper-sensitivity. Approximately 10^7 cells were serially diluted 10-fold on the above-mentioned plates and were incubated at 30°C for 24-48 hours before imaging. Experimental replicates, n = 3. (2X) indicates the cells spotted approximately double in number to compensate slow growth rate. **(B)** Budding index of strains indicated in (A) under standard growth conditions. N = 200, Experimental replicates, n = 2 **(C)** Left, representative images of the live cells of the strains indicated in (A) showing segregation of chromatin (DAPI) and SPBs (Spc42-EGFP). Scale bar = 2 µm. In the ‘defective segregation’ panel, the double-headed arrow denotes the distribution of DAPI mass away from the bud-neck. Right, the percentage of large-budded cells with ‘normal’ and ‘defective’ segregation of DAPI/SPBs was shown (with cartoons) for the indicated strains. N = 200, Experimental replicates, n = 2. **(D)** Time-lapse imaging of live cells (see materials and methods) to visualize kinetochore (Ndc80-CFP) and SPB (Spc42-EGFP) dynamics in the indicated strains. The small budded cells with single SPB and kinetochore signals (at S-Phase) were considered as the start-point (t = 0 min) for both wild-type and mutant cells, N = 12. The arrowheads denote the formation of kinetochore/SPB bi-lobed configuration. Scale bar = 2 µm.

Since the benomyl sensitivity (Fig 1A) and the observed slow growth of the *bmh* mutants are indicative of defects in chromosome segregation, we wished to know at which stage of the cell cycle the defect is occurring. The budding index revealed that a high percentage of large-budded cells accumulated in *bmh1Δ bmh2Δ* double mutant (Fig 1B) indicating cells are being delayed at G2/M and/or at anaphase. To pinpoint the cell cycle stage of the delay, we analysed the segregation pattern of the DNA mass (DAPI) and spindle pole bodies/SPBs (Spc42-EGFP) in the large-budded cells (Fig 1C). We found that while the wild-type and the single mutants showed normal segregation (equally segregated DAPI/SPBs signal in mother and daughter buds), the *bmh1Δ bmh2Δ* double mutant largely (∼60%) showed defective segregation (undivided single DAPI mass with both SPBs in the mother bud). Notably, in the cells showing defective segregation, the presence of DAPI and the spindle well away from the bud neck (double-headed arrow) despite having a large bud indicates that the movement of the nucleus to the bud neck and subsequent separation of the SPBs are perhaps delayed in *bmh1Δ bmh2Δ* double mutant. In order to visualize the dynamics of kinetochore and SPB separation, we fluorescently labelled kinetochores (Ndc80-CFP) and SPBs (Spc42-GFP) in the wild-type and *bmh1Δ bmh2Δ* double mutant and observed the cell cycle by time-lapse imaging of the live cells (Fig 1D). Considering that the cells are in similar cell cycle stage at the start point (t = 0 mins) (see material and methods), a delay in the successful duplication followed by the bi-lobed metaphase configuration of kinetochores and SPBs (marked by arrowhead) was observed in *bmh1Δ bmh2Δ* double mutant (t = 60 mins) compared to the wild-type (t = 20 mins). The delay in forming the bi-lobed configuration in *bmh1Δ bmh2Δ* mutants may be because of previously reported G1/S transition delay (Lottersberger et al., 2006). After forming the bi-lobed configuration, the wild-type cells successfully segregated their kinetochores and SPBs between mother and daughter buds within the range of normal doubling time (t = 80-100 mins); in contrast, *bmh1Δ bmh2Δ* double mutant displayed a substantial delay in further segregation where the SPBs remained within the mother bud even after 180 min indicating cells were either delayed at metaphase to anaphase transition or even after anaphase was triggered, SPBs were not able to separate due to physical constraints. Interestingly, compared to the wild-type, a reduced intensity of the kinetochore (Ndc80-CFP) signal was consistently observed in *bmh1Δ bmh2Δ* double mutant after achieving a bi-lobed configuration (t = 60 mins). The observed reduction in the kinetochore signal may be attributed to potential kinetochore perturbation. We conclude that loss of both the Bmh proteins cause a high percentage of chromosome missegregation perhaps due to defects in KT-MT related functions. However, negligible growth defects (Fig 1B, C) in the corresponding *bmh* single mutants imply occurrence of functional complementation between the Bmh proteins despite their independent contribution to proper chromosome segregation as per earlier report (Wilkert et al., 2005).

### Bmh proteins are necessary for stable KT-MT attachment dynamics during metaphase

Since *bmh1Δ bmh2Δ* double mutant cells showed sensitivity to the anti-microtubule drug (Fig 1A), loss of kinetochore signal after formation of bi-lobed configuration (Fig 1D), and a substantial delay in anaphase separation of the chromosomes (Fig 1D), we hypothesize that perhaps the KT-MT attachment dynamics are perturbed in these cells. Earlier studies have reported that non-essential kinetochore proteins contribute to proper organization of the kinetochore and thereby promote high-fidelity chromosome segregation (reviewed in Mehta et al 2022). Therefore, we wished to examine the organization of the kinetochore in the absence of Bmh proteins using chromatin spread with a hypothesis that localization of the kinetochore proteins at the centromeres will be altered without the Bmh proteins. Since we observed an accumulation of *bmh1Δ bmh2Δ* cells with undivided DAPI (Fig 1C), we wished to compare the localization of kinetochore proteins in the cells with undivided DAPI mass. On the chromatin spreads made from asynchronously grown wild-type and the *bmh1Δ bmh2Δ* double mutant cells we chose to monitor the localization of the epitope-tagged outer kinetochore protein, Ndc80-HA, which was used previously (Janke et al., 2001). Based on the Ndc80-HA signal within undivided DAPI mass, we categorized the spreads as follows: ‘clustered kinetochores’, ‘declustered kinetochores’, and ‘weak-signal’ represent either single or bi-lobed tight-knit foci, multiple foci and multiple weak intensity foci, respectively (Fig 2A). In comparison with wild-type, *bmh1Δ bmh2Δ* double mutants displayed a significant increase in the percentages of spreads harboring declustered kinetochores or weak-signal (∼57%) with nearly a cumulative equivalent percentage was reduced in the clustered kinetochores category (Fig 2B). The observed increase in the percentage of spreads exhibiting weakened signals and declustered kinetochores implies two different contributions of Bmh proteins viz., they may play a crucial role in stabilizing the kinetochores and in keeping them clustered. We further biochemically assessed the localization of Ndc80-HA at the centromeres by chromatin immunoprecipitation (ChIP) assay using anti-HA antibodies in asynchronously grown wild-type and *bmh1Δ bmh2Δ* cells. In coherence with chromatin spread, a reduced binding of Ndc80-HA was observed at the centromeres in *bmh1Δ bmh2Δ* double mutant compared to the wild-type (Fig 2C) indicating the contribution of Bmh proteins in kinetochore stability. To explore the role of Bmh proteins in kinetochore clustering, we fluorescently labelled kinetochores (Ndc80-CFP) and SPBs (Spc42-EGFP) and performed live-cell imaging to compare kinetochore distribution between wild-type and *bmh1Δ bmh2Δ* double mutant during metaphase. We determined the cell cycle stage of the cells based on bud size and SPB-SPB distance, as previously defined (Tytell and Sorger, 2006). Based on the Ndc80-CFP signal, we categorized the metaphase cells into ‘clustered kinetochores’ showing tight-knit bi-lobed foci between two SPBs and ‘declustered kinetochores’ showing multiple foci linearly distributed between two SPBs (Fig 2D, left). We observed a substantial percentage of cells with declustered kinetochores in *bmh1Δ bmh2Δ* double mutant (>70%) as compared to the wild-type (∼20%) (Fig 2D, right). For better visualization of the distribution of the kinetochores (Ndc80-CFP) line scans were performed between SPBs which clearly depicted deterioration of bimodal distribution of the kinetochore clusters in the double mutant (Fig 2E, bottom) compared to the wild-type (Fig 2E, top). The observed defect in kinetochore protein localization and in its clustering is not specific to Ndc80, as a similar pattern was observed for another kinetochore protein, Mtw1 in *bmh1Δ bmh2Δ* double mutant (Fig S2A-E). Additionally, to rule out the possibility of differences in the overall expression level of kinetochore proteins in the absence of Bmh proteins, we estimated and compared protein levels of Ndc80 and Mtw1 in the wild-type and *bmh1Δ bmh2Δ* double mutants by western blotting and found no significant difference (Fig S3A, B). Taken together, we conclude that in absence of both the Bmh proteins the kinetochore integrity is compromised which perhaps alters the dynamics of KT-MT attachment causing kinetochores to be irregularly distributed (declustered) between two SPBs.

**Figure 2:**
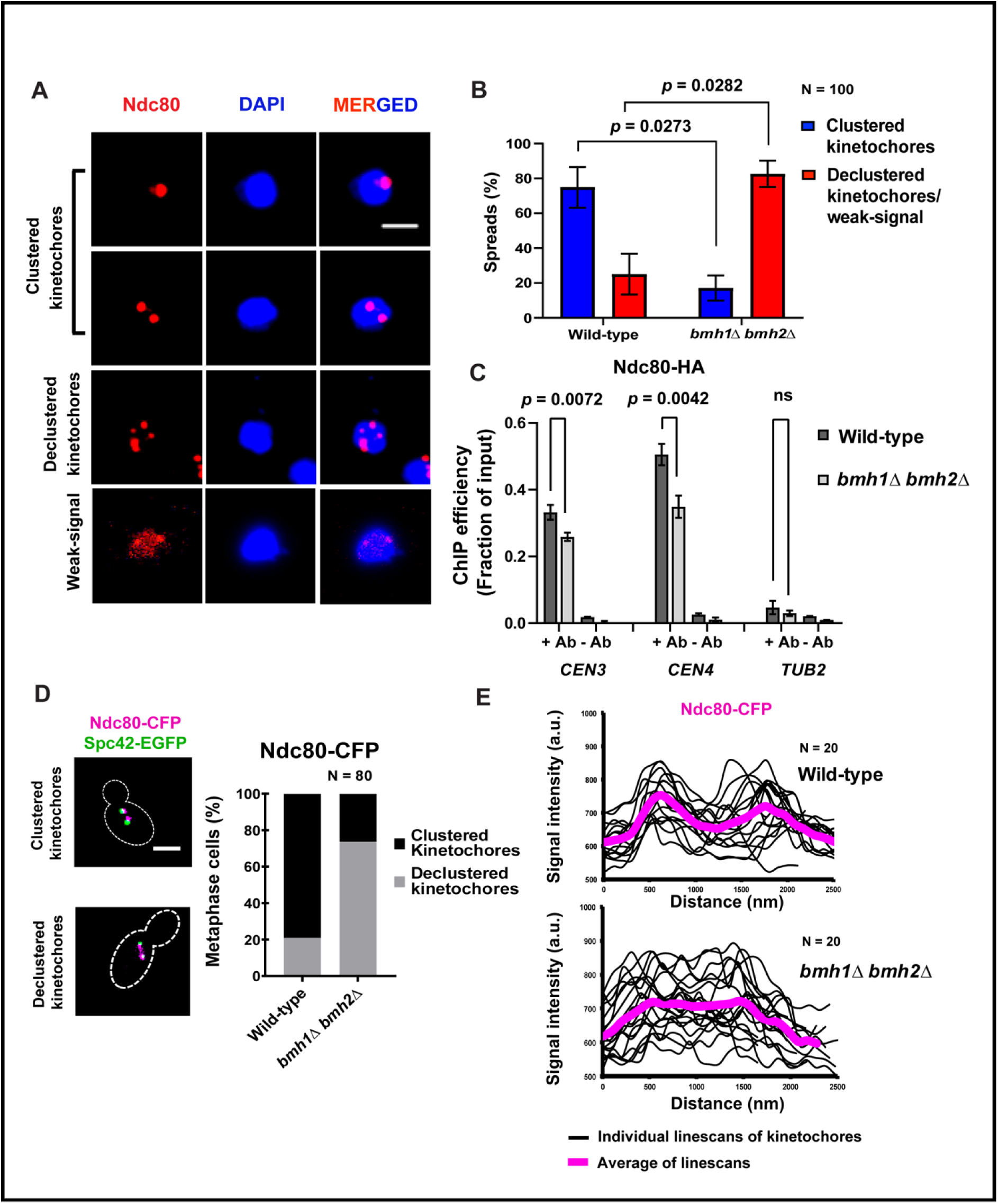
Perturbed clustering and integrity of the kinetochores (Ndc80) in the absence of Bmh proteins. (**A**) The representative images of the chromatin spreads to visualize kinetochore protein Ndc80 fused to HA in wild-type and *bmh1Δ bmh2Δ* cells before anaphase (undivided DAPI mass). Anti-HA antibodies and DAPI staining were used to probe Ndc80-HA and chromatin, respectively. Scale bar = 2 µm. **(B)** The percentage of different categories of the spreads as mentioned in A, for the indicated strains. N = 100, Experimental replicates, n = 2. **(C)** ChIP-qPCR analyses were performed to quantify the association of Ndc80-HA with *CEN3, CEN4,* and *TUB2* (negative control) loci in the indicated strains. Anti-HA antibodies were used for pull-down from the asynchronously grown mid-log cells. Error bars indicate standard error. Experimental replicates, n = 3. Statistical significance (*p*) was calculated using the two-tailed student’s t-test. ‘ns’ represents statistically not significant. **(D)** Left, representative live cell images showing localization pattern of Ndc80-CFP during metaphase (SPB-SPB distance = 1.5 – 2.5 μm). Scale bar = 2 µm. Right, the percentage of metaphase cells depicting ‘clustered’ or ‘declustered’ kinetochores were shown for the indicated strains. N = 100, Experimental replicates, n = 2. (E) The intensity graphs depict the distribution of fluorescence signal intensity of Ndc80-CFP (Magenta) in the indicated strains. Line scans were performed along the axis between two SPBs for 20 cells to estimate mean signal intensity distribution (arbitrary units) for each strain. Experimental replicates, n = 2.

Alternatively, the kinetochore declustering can happen due to defective spindle formation in absence of the Bmh proteins. To test this, we observed spindle morphology along with kinetochore (Mtw1) localization in wild-type and *bmh1Δ bmh2Δ* double mutants. Irrespective of the kinetochore conformation (clustered or declustered), no visible difference in the spindle morphology was observed (Fig S3C, D). In support of our observation, unperturbed spindle morphology in cells exhibiting declustered kinetochores was reported earlier in kinesin motor mutants (Tytell and Sorger, 2006). In conclusion, our findings suggest that Bmh proteins are essential for stable KT-MT attachment dynamics, majorly through the regulation of kinetochore, but not spindle, functions.

### Bmh proteins are necessary for chromosome congression during metaphase

The formation of bi-lobed clusters of sister kinetochores requires synchronized poleward movement of the chromosomes, termed as chromosome congression, following their attachments to the kinetochore microtubules (kMTs) (Xiangwei et al., 2000; Pearson et al., 2004). Any defect in chromosome congression can be identified by the following observations. First, Hyperstretching of the pericentromeric chromatin due to deregulated pulling/pushing forces resulting from alteration of plus– and/or minus-end growth-shrinkage dynamics of the kMTs during metaphase-anaphase transition (Thrower and Bloom, 2001; Tytell and Sorger, 2006; Wargacki et al., 2010). Second, an accumulation of lagging chromosomes results in a linearly declustered distribution of the kinetochores between two SPBs (Tytell and Sorger, 2006). To examine if chromatin hyperstretching is occurring, we fluorescently labelled the pericentric chromatin at 1.4 kb away from *CEN5* using TetO/TetR-GFP system (Michaelis et al., 1997) in the wild-type and *bmh1Δ bmh2Δ* cells harboring fluorescently labelled SPBs (Spc42-mCherry). We selectively scored cells either in the metaphase or in early anaphase stage judged by the SPB-SPB distance (2.0 – 4.0 μm) and categorized the behaviour of the sister chromatids based on the type of *CEN5*-GFP fluorescent signal as follows (Fig 3A). Type-I and Type-II, two sister *CEN5*-GFPs are coalesced into one focus or divided into two foci, respectively; Type-III, cells with either one (top row) or two (bottom row) hyperstretched *CEN5*-GFP signals. Here, type-I represents the cells in early metaphase stage (SPB-SPB distance < 2.5 μm), whereas type-II and – III represent the cells in late metaphase to early anaphase cells (SPB-SPB distance = 2.5 – 4.0 μm). While comparing with wild-type, we observed a significant increase in the percentage of type-III population (∼27%) in *bmh1Δ bmh2Δ* double mutants; while nearly an equivalent percentage was reduced in type-II population (∼24%), and no significant change in type-I population (∼3%) was observed (Fig 3B). These data indicate that the cells with hyperstretched chromatids are getting accumulated in *bmh1Δ bmh2Δ* double mutants specifically during the metaphase-anaphase transition and the hyperstretching is not prevalent during the early metaphase stages. To be noted, pericentric hyperstretching denotes a strong poleward force exerted over centromeres by the kMTs, which is possible only after the successful establishment of KT-MT attachment. Since nearly 50% of the *bmh1Δ bmh2Δ* cells show hyperstretched chromatin that indicates KT-MT attachment is not perturbed in majority of the cells.

**Figure 3:**
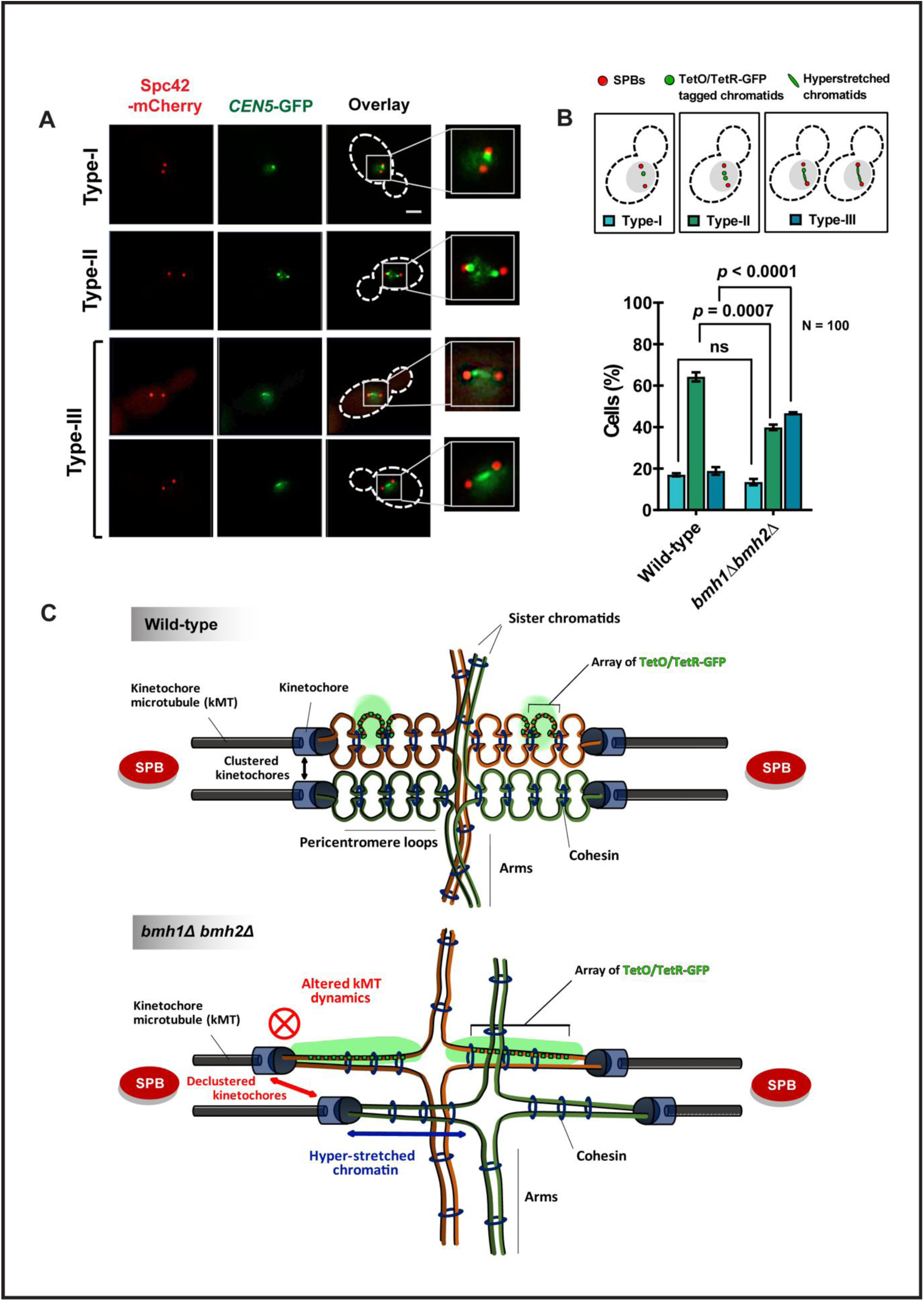
Chromatin hyperstretching near centromeres in *bmh* double mutants. (**A**) Representative images of the live cells depicting the status of the chromatin proximal to *CEN5* marked by TetO/TetR-GFP in the indicated strains during metaphase to anaphase transition. Scale bar = 2 µm. **(B)** The percentage of the cells of ‘type-I – III categories (with cartoons) as shown in (A), are graphically represented for the indicated strains. Error bars indicate standard error. N = 90 – 120, Experimental replicates, n = 3. Statistical significance (*p*) was calculated using the two-tailed student’s t-test. ‘ns’ represents statistically not significant. **(C)** Schematics of kinetochore and pericentric chromatin behavior in wild-type and *bmh1Δ bmh2Δ* mutant during the metaphase-anaphase transition. Top, in wild-type cells, the kinetochores stabilize the kMT dynamics and thereby maintain the clustering of the kinetochores from different chromosomes (black double-headed arrow) to form a bi-lobe cluster architecture during metaphase. Consequently, the signals from the array of TetO/TetR-GFP (green boxes) integrated within the pericentromeric loop of each sister chromatid coalesce into a single focus (green blob). Bottom, in *bmh1Δ bmh2Δ* cells, perturbed kinetochores fail to maintain uniform kMT dynamics, resulting in random positioning of the kinetochores (red double-headed arrow) along the spindle axis causing the linearly declustered distribution. Altered kMT dynamics also cause hyperstretching of the peri-centromeric loops (blue double-headed arrow) in which the signal from TetO/TetR-GFP arrays appears elongated (green stretched signal).

Previously observed linear declustering of the kinetochores (Fig 2D) clearly demonstrates the presence of lagging chromosomes between two SPBs during metaphase. However, similar measurements in anaphase cells (SPB-SPB distance > 4 μm) showed no significant increase in declustering phenotype in the mutant over the wild-type suggesting an amelioration of the kinetochore clustering with time (Fig S4A, B). In coherence with our observation, the cells with defective kinesin-8 motor (*kip3Δ*) displayed lagging chromosomes with a delay in the duration of chromatid hyperstretching near centromeres due to altered microtubule dynamics during the metaphase-anaphase transition (Tytell and Sorger, 2006; Wargacki et al., 2010). Overall, we conclude that the kinetochore declustering (Fig 2A, D) and chromatin hyperstretching (Fig 3A, B) phenotypes observed in *bmh1Δ bmh2Δ* double mutants are perhaps due to defective chromosome congression which is a consequence of the altered kMT dynamics; as shown by the schematics (Fig 3C).

### The G2/M delay in *bmh1Δ bmh2Δ* double mutant is Mad2 dependent

It has been shown earlier that mutants with perturbations in KT-MT attachment related functions activate SAC and are synthetically lethal or sick with the SAC mutants (Wang and Burke, 1995; Pangilinan and Spencer, 1996; Hyland et al., 1999; Cheeseman et al., 2001; Daniel et al., 2006). As *bmh1Δ bmh2Δ* cells displayed both kinetochore perturbations and defective kMT dynamics, we believed that the observed G2/M delay (Fig 1B, D) is SAC dependent and therefore *bmh1Δ bmh2Δ* double mutant is expected to interact genetically with the SAC mutants. To examine this, *bmh1Δ bmh2Δ* mutant was crossed with an isogenic strain deleted for *MAD2*, the major SAC component (Li and Murray, 1991). The resulting diploids were sporulated and tetrads were dissected. We observed that the *mad2Δ bmh1Δ bmh2Δ* triple mutant spores were viable (Fig 4A, solid square) but their growth was weaker than the *bmh1Δ bmh2Δ* double mutant spores (Fig 4A, dotted squares) indicating a synthetic negative interaction between *mad2Δ* and *bmh1Δ bmh2Δ*. Consequently, in presence of a sub-lethal concentration of benomyl *mad2Δ bmh1Δ bmh2Δ* cells performed very poorly compared to the corresponding double or single mutants (Fig 4B). Notably, *mad2Δ bmh1Δ* double mutant grew poorer than the *mad2Δ bmh2Δ*. This is not unexpected as Bmh1 has been shown previously to be involved in SAC silencing and SPOC checkpoint function (Grandin and Charbonneau, 2008; Caydasi et al., 2014). Interestingly, the *mad2Δ bmh1Δ bmh2Δ* cells grew poorer than the *mad2Δ bmh1Δ* cells, indicating that Bmh2 contributes towards kinetochore related functions unshared with Bmh1.

**Figure 4:**
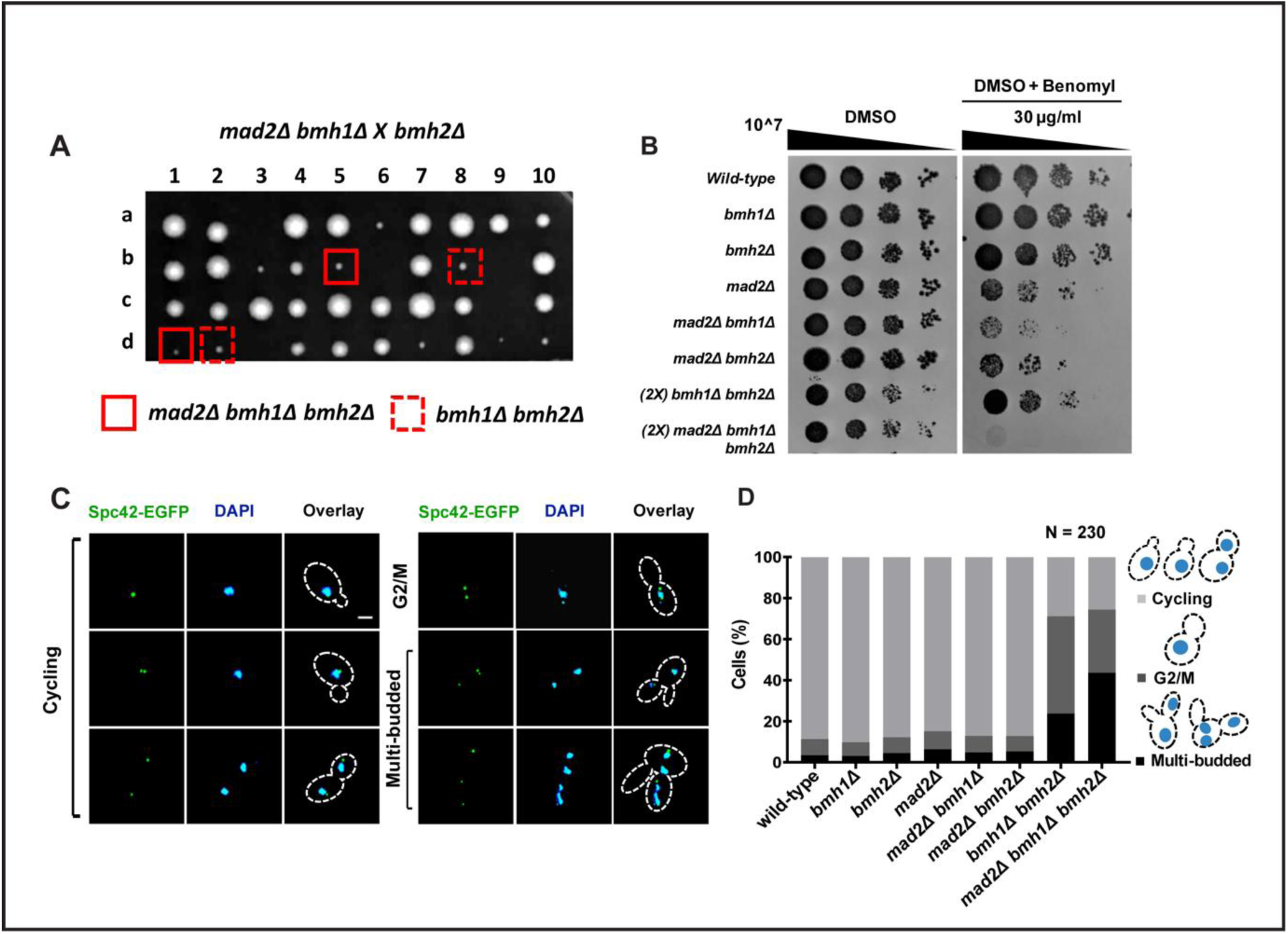
Mad2 mediated G2/M delay in *bmh* double mutants. (**A**) Genetic interactions of *bmh* mutants with a SAC mutant, *mad2Δ* was assessed by the growth of the spores harboring double (*mad2Δ bmh1Δ, mad2Δ bmh2Δ, bmh1Δ bmh2Δ*) or triple mutations (*mad2Δ bmh1Δ bmh2Δ*). The triple mutant was obtained by crossing *mad2Δ bmh1Δ* and *bmh2Δ* mutants, whereas the double mutants were obtained by crossing appropriate single mutants. The plates were incubated at 30°C for 3-5 days before they were photographed. **(B)** The depicted wild-type, single, double, and triple mutants were spotted on control (DMSO) and indicated concentrations of drug (DMSO+Benomyl) containing YEPD plates. Approximately 10^7 cells were spotted following 10-fold serial dilution on the above-mentioned plates and were incubated at 30°C for 48-54 hours before imaging. Extended incubation time was provided for the recovery of hypersensitive strains to highlight subtle differences. Experimental replicates, n = 3. (2X), indicates the cells spotted approximately double in number to compensate slow growth rate. **(C)** The representative live cell images of the cycling, G2/M, and multi-budded cells based on their bud morphology and distribution of chromatin (DAPI) and SPBs (Spc42-EGFP) within mother and daughter cell compartments. Scale bar = 2 µm. **(D)** The percentages of cells under indicated categories are shown (with cartoons) for the indicated strains grown under standard conditions. Error bars indicate standard error. N = 230, Experimental replicates, n = 3.

Abrogation of SAC-mediated G2/M arrest in presence of any perturbations in KT-MT interaction leads to progression into the cell cycle that generates multi-bud phenotype in budding yeast (Grandin and Charbonneau, 2008; Caydasi et al., 2014). Since we observed that the *bmh* mutants genetically interact with *mad2Δ* (Fig 4A, B) indicating that Bmh proteins may also have a role in SAC activation, we hypothesize that a fraction of *bmh* mutant cells may abrogate SAC-mediated G2/M arrest and generate multi-budded cells. While assessing phenotypes of the budded cells, we broadly categorized them as ‘cycling’ (small-budded cells with undivided DAPI with two or one SPB(s) in the mother compartment plus the large-budded cell with one each of DAPI and SPB in both the compartments), ‘G2/M’ (large-budded cells harboring undivided DAPI with two SPBs within mother compartment), and ‘multi-budded’ (cells with multiple buds containing fragmented DAPI mass and super-numerary SPBs). We indeed observed a significant fraction of cells (∼23%) showed multi-bud phenotype in *bmh1Δ bmh2Δ* double mutant indicating that although there is a sizable fraction (∼47%) of cells at G2/M stage accounting for SAC mediated arrest, the double mutant is partially compromised in that arrest (Fig 4D). Further removal of Mad2 in these cells caused, as expected, a nearly two-fold increase (∼44%) in the multi-bud phenotype (Fig 4D, compare black bars of *bmh1Δ bmh2Δ* and *mad2Δ bmh1Δ bmh2Δ* mutants) with a nearly equivalent reduction in the percentage of G2/M arrest phenotype again indicating that the latter phenotype in *bmh1Δ bmh2Δ* is largely dependent on SAC. However, compared with the wild-type (∼4%), no statistically significant increase in the percentage of multi-budded phenotype was observed in *mad2Δ bmh1Δ* (∼8%) or *mad2Δ bmh2Δ* (∼7%) cells, perhaps due to functional complementation between the Bmh proteins. Altogether, we conclude that the G2/M delay observed in the *bmh1Δ bmh2Δ* double mutants (Fig 1B, D) is perhaps due to Mad2-dependent transient activation of the SAC in response to the altered KT-MT attachment dynamics.

### Bmh proteins contribute to kinetochore function

Since *bmh1Δ bmh2Δ* double mutant cells showed sensitivity to the anti-microtubule drug and the cells showed a delay in anaphase separation of the chromosomes (Fig 1A, D), we hypothesize that perhaps the kinetochore function is weak in these cells. This is further fuelled by the facts that we could identify strong Bmh protein recognition motifs in multiple kinetochore proteins (Kumar, 2018), and others have reported a physical interaction between the kinetochore protein Iml3 and Bmh1-binding protein Fin1 (Kakiuchi et al., 2007; Akiyoshi et al., 2009). Therefore, we utilized W303 background strain (in which *bmh* double mutant is lethal) to explore the distinct functions of individual Bmh proteins and to eliminate the possibility of alternative cellular pathways (later mentioned in Fig 7) to functionally complement the loss of the Bmh proteins. To assess this, we first examined if there is any genetic interaction between *bmh* and kinetochore mutants. For this, we used a non-essential kinetochore protein mutant, *ctf19Δ* (Hyland et al., 1999). We could not find any synthetic growth defect in *ctf19Δ bmh1Δ* or *ctf19Δ bmh2Δ* double mutants under normal conditions (Fig 5A, DMSO). However, in presence of a sub-lethal concentration of benomyl, where *ctf19Δ* showed considerable growth defect as reported earlier (Hyland et al., 1999), *ctf19Δ bmh2Δ* double mutant grew worse than *ctf19Δ* (Fig 5A, DMSO + Benomyl; compare *ctf19Δ* with *ctf19Δ bmh2Δ*). Interestingly, the growth of *ctf19Δ* became marginally better upon removal of Bmh1 (Fig 5A, compare *ctf19Δ* with *ctf19Δ bmh1Δ*). These results suggest that the loss of Bmh proteins, Bmh2 in particular, might influence the kinetochore function.

**Figure 5:**
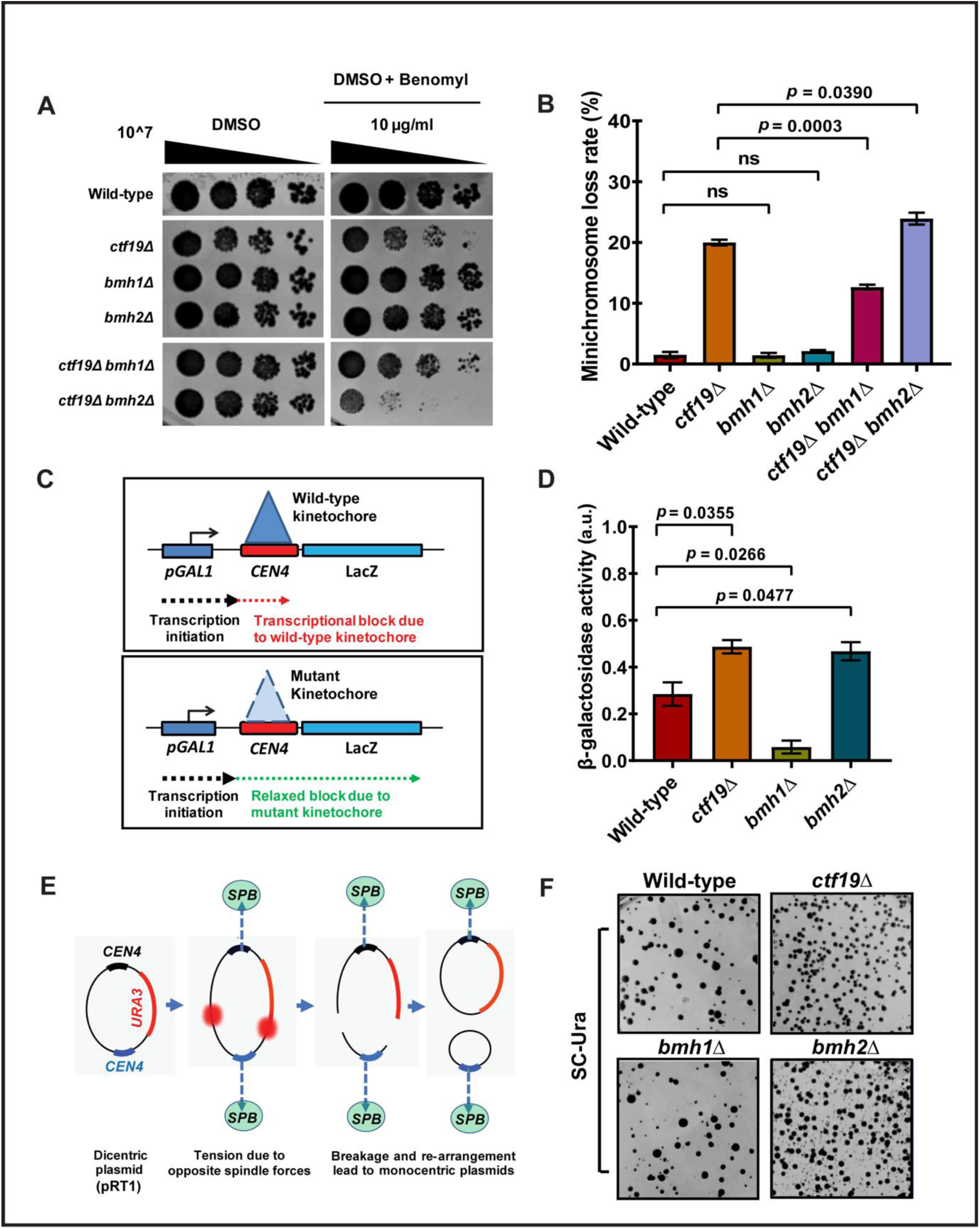
Bmh proteins individually contribute to the integrity of the kinetochore ensemble. (**A**) Genetic interactions of *bmh* mutants with a kinetochore mutant, *ctf19Δ*. Wild-type, single (*ctf19Δ, bmh1Δ, bmh2Δ),* and double (*ctf19Δ bmh1Δ, ctf19Δ bmh2Δ*) mutant cells were spotted on control (DMSO) and indicated concentrations of drug (DMSO+Benomyl) containing YEPD plates. Approximately 10^7 cells were serially diluted 10-fold on the above-mentioned plates and were incubated at 30°C for 24-48 hours before imaging. Experimental replicates, n = 5 **(B)** Minichromosome (YCplac33 plasmid) loss rate was estimated (see materials and methods) in the strains used in (A) from three independent transformants from each strain. Error bars indicate standard error. Experimental replicates, n = 3. Statistical significance (*p*) was calculated using the two-tailed student’s t-test. ‘ns’ represents statistically not significant. **(C)** The schematic of the transcriptional read-through assay to probe the integrity of the kinetochore complex by scoring the level of transcription across kinetochore formed on a centromere (*CEN*) placed between *GAL1* promoter and LacZ reporter gene. **(D)** The β-galactosidase activity from the wild-type, *ctf19Δ, bmh1Δ*, and *bmh2Δ*, cells harboring integrated *pGAL1*-*CEN4*-LacZ cassette as described in materials and methods. Error bars indicate standard error. Experimental replicates, n = 3. Statistical significance (*p*) was calculated using the two-tailed student’s t-test. **(E)** The schematic of a di-centric plasmid harboring *URA3* genetic marker and its re-arrangement to generate monocentric plasmids in a mitotic cell. The round red shapes indicate the random breakage points **(F)** The mitotic stability of the dicentric plasmid shown in (E), in wild-type, *ctf19Δ*, *bmh1Δ*, and *bmh2Δ* cells was qualitatively estimated by the presence of either homogeneity or heterogeneity in the colony size on the indicated plate. Experimental replicates, n = 3. The strains used above in (A-F) are from W303 genetic background strains, in which *bmh* double mutant is inviable.

The requirement of any non-essential protein for fine tuning of functional kinetochore in budding yeast can be assessed by measuring the mitotic stability of minichromosomes (Maine et al, 1984; Sanyal et al 1998, Poddar et al 1999, Ghosh et al 2001) and by directly determining the kinetochore structural integrity using transcriptional read-through assay and dicentric stability assay (Mann and Davis, 1983; Koshland et al., 1987, Doheny et al., 1993). Coherent with the previous observation (Fig 5A), the enhanced loss of the minichromosome in *ctf19Δ bmh2Δ* double mutant compared to *ctf19Δ* or *bmh2Δ* single mutant clearly denotes the requirement of Bmh2 for the proper function of the kinetochore (Fig 5B). Conversely, consistent with the results of synthetic interactions (Fig 5A), removal of Bmh1 in *ctf19Δ* cells rescued the minichromosome loss phenotype to some extent suggesting that loss of Bmh1 may positively influence the kinetochore function.

In the transcriptional read-through assay, transcription across a centromere and a reporter placed downstream of the centromere is perturbed in the wild-type due to kinetochore complex formation at the centromere (Doheny et al., 1993). However, in cells with compromised kinetochore complex, the transcription can proceed causing the reporter expression (Fig 5C). The β-galactosidase activity, as a read-out of LacZ reporter expression, from the *bmh* mutants and kinetochore mutant *ctf19Δ* was compared with the wild-type. Similar to the *ctf19Δ* mutant, the relative β-galactosidase activity was found high in *bmh2Δ* but not in *bmh1Δ* mutant (Fig 5D), indicating the requirement of Bmh2 but not Bmh1 for kinetochore structural integrity. Similarly, the dicentric plasmid stability assay was also used to judge the essentiality of a protein to the kinetochore function in yeast cells (Mann and Davis, 1983; Koshland et al., 1987). The dicentric plasmids within wild-type cells end up with breakage due to microtubule-mediated opposite pulling force acting on two kinetochores formed on two centromeres. Hence, a dicentric plasmid will be unstable and will be converted into monocentric plasmid of different sizes in different cell lineages upon rearrangements and ligation (Fig 5E). Consequently, the colonies that appeared after transformation of the cells with dicentric plasmid will be of heterogeneous sizes (Koshland et al., 1987; Doheny et al., 1993; Sanyal et al., 1998). Conversely, a kinetochore mutant cell due to weak KT-MT interaction avoids the breakage causing the dicentric plasmid to be stabilized, producing homogenous transformant colonies. As expected from the above or earlier reports, *ctf19Δ* and *bmh2Δ* mutants produced homogenous colonies indicating harboring compromised kinetochore integrity, whereas *bmh1Δ* mutant showed heterogenous colonies like in the wild-type (Fig 5F).

In summary, the above observations together suggest that Bmh proteins are important for proper functioning of budding yeast kinetochore. Among the Bmh proteins, Bmh2 promotes the functions of the kinetochore perhaps by influencing its integrity. On the other hand, a significant increase in mitotic stability of a minichromosome in *ctf19Δ bmh1Δ* over *ctf19Δ* alone (Fig 5B), β-galactosidase activity lower than the wild-type in transcriptional read-through assay (Fig 5D), and dicentric plasmid instability (Fig 5F) in *bmh1Δ* hint towards a positive impact of loss-of-Bmh1 on kinetochore integrity. As a corollary, overall stress tolerance and enhancement in the life span of budding yeast cells with *bmh1Δ* mutation were reported earlier (Wang et al., 2009).

### Bmh proteins partially co-localize with kinetochore throughout the mitosis

Bmh-interacting motifs have been identified in several kinetochore and SPB proteins (Kumar, 2018); and in a previous high-throughput screening of direct physical interactors of Bmh proteins, kinetochore protein Iml3 was co-purified (Kakiuchi et al., 2007). These observations and our findings strongly suggest that Bmh proteins contribute to the kinetochore function. In this context, we hypothesize that Bmh proteins physically present at the kinetochores to execute their roles. To investigate this, we constructed strains where both Ndc80 and either Bmh1 or Bmh2 were tagged with 6XHA and 13XMyc at their C-terminals, respectively. The fused proteins were functional as evident from earlier studies (Janke et al., 2001; Slubowski et al., 2014). Their localizations were assessed simultaneously on chromatin spreads made from the cells harvested from different stages of the cell cycle that were judged from the spreads as follows: G1/S or metaphase stage as single DAPI mass harboring a single Ndc80 focus (Fig 6A) or bi-lobed Ndc80 foci (Fig 6B), respectively; anaphase stage as two (separated or connected) DAPI masses each harboring one Ndc80 focus (Fig 6C). Since Bmh proteins are known for their binding to the cruciform DNA and they localize on the chromatin at the autonomously replicating sequence (*ARS*) sites (Yahyaoui et al., 2007; Yahyaoui and Zannis-Hadjopoulos, 2009), we expected multiple foci of Bmh proteins of which a subset will be kinetochore-specific on the spreads. We indeed observed chromatin localization of Bmh proteins as multiple foci on the spreads from different cell cycle stages (Fig 6A-C). From each cell-cycle stage, the extent of co-localization of Ndc80 and Bmh1/2 was measured using Pearson’s correlation coefficient (PCC), and the obtained values were compared with no-tag control for statistical significance (Fig 6D). After thorough analysis (materials and methods), we observed a significant percentage (> 60%) of Bmh proteins co-localize either partially or completely with Ndc80 throughout the cell cycle (Fig 6E). The co-localization patterns ‘No’ (PCC < 0.1), ‘partial’ (PCC = 0.1-0.5), and ‘complete’ (PCC > 0.5) were categorized based on PCC values as described elsewhere (Zinchuk and Grossenbacher-Zinchuk, 2014; Prajapati et al., 2017). These results indicate that at least a fraction of Bmh proteins may reside at or close to the centromeres.

**Figure 6:**
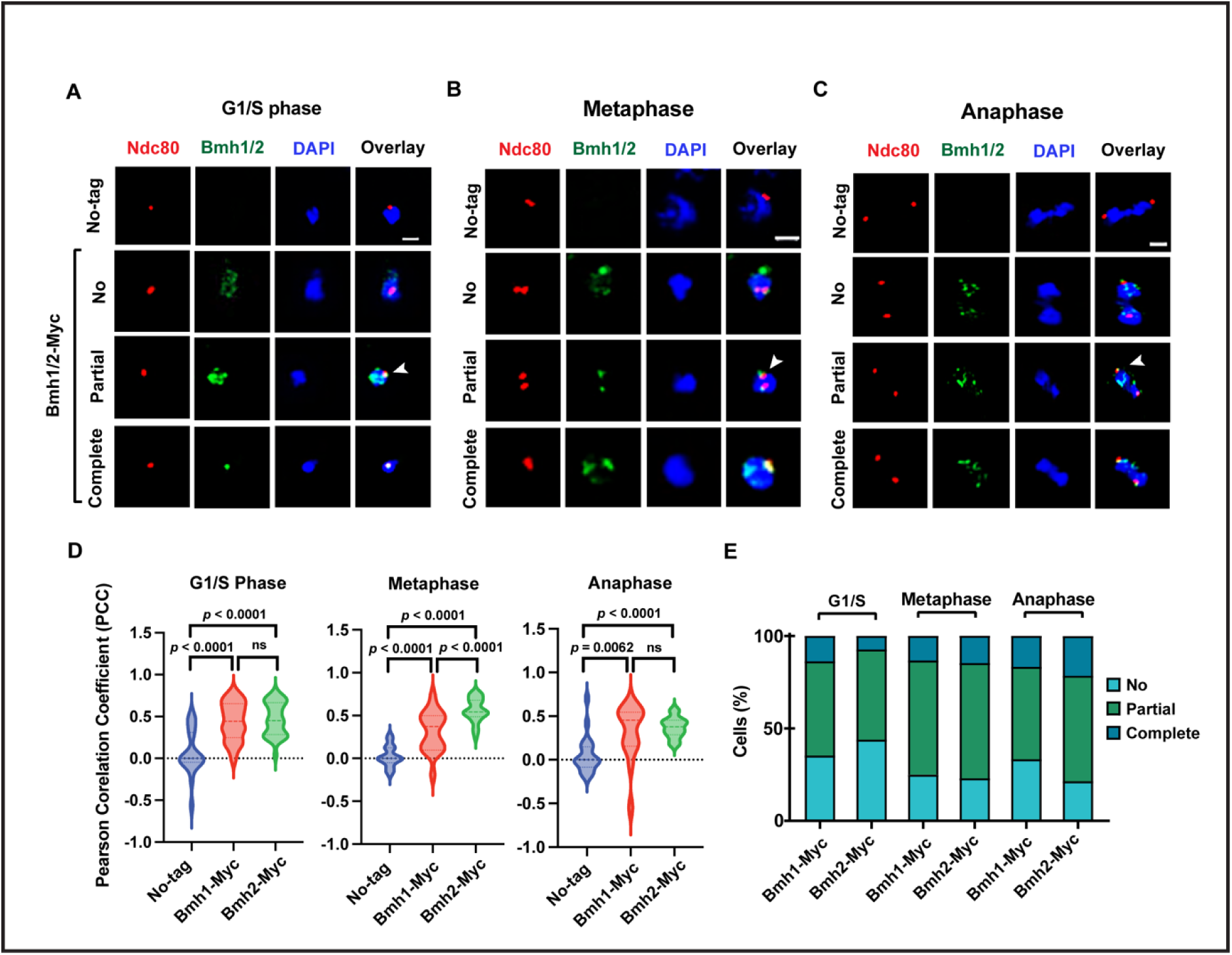
Co-localization of Bmh proteins with outer kinetochore protein Ndc80. The representative images of the chromatin spreads to visualize co-localization of Bmh proteins with kinetochore protein Ndc80 during **(A)** G1/S, **(B)** metaphase, and **(C)** anaphase cell cycle stages. Anti-Myc, anti-HA, and DAPI staining were used to probe Bmh proteins (Bmh1/2-Myc), kinetochore (Ndc80-HA), and chromatin, respectively, in no-tag (Ndc80-HA), Bmh1-Myc (Bmh1-Myc Ndc80-HA), and Bmh2-Myc (Bmh2-Myc Ndc80-HA) strains. The extent of colocalization of Ndc80 and Bmh proteins is categorized as ‘no’, ‘partial’ (arrowhead), and ‘complete’. Scale bar = 2 µm. **(D)** Quantification of the colocalization between Ndc80 and Bmh1/Bmh2 using Pearson’s correlation coefficient (PCC). The estimated PCC values from Bmh1/2-Myc strains were compared with no-tag strains to derive statistical significance. Error bars indicate standard error. N = 80, Experimental replicates, n = 2. Statistical significance (*p*) was calculated using the two-tailed student’s t-test. ‘ns’ represents statistically not significant. **(E)** The percentage of the spreads with different co-localization categories from the cells of indicated strains and cell-cycle stages. N = 80, Experimental replicates, n = 2.

**Figure 7:**
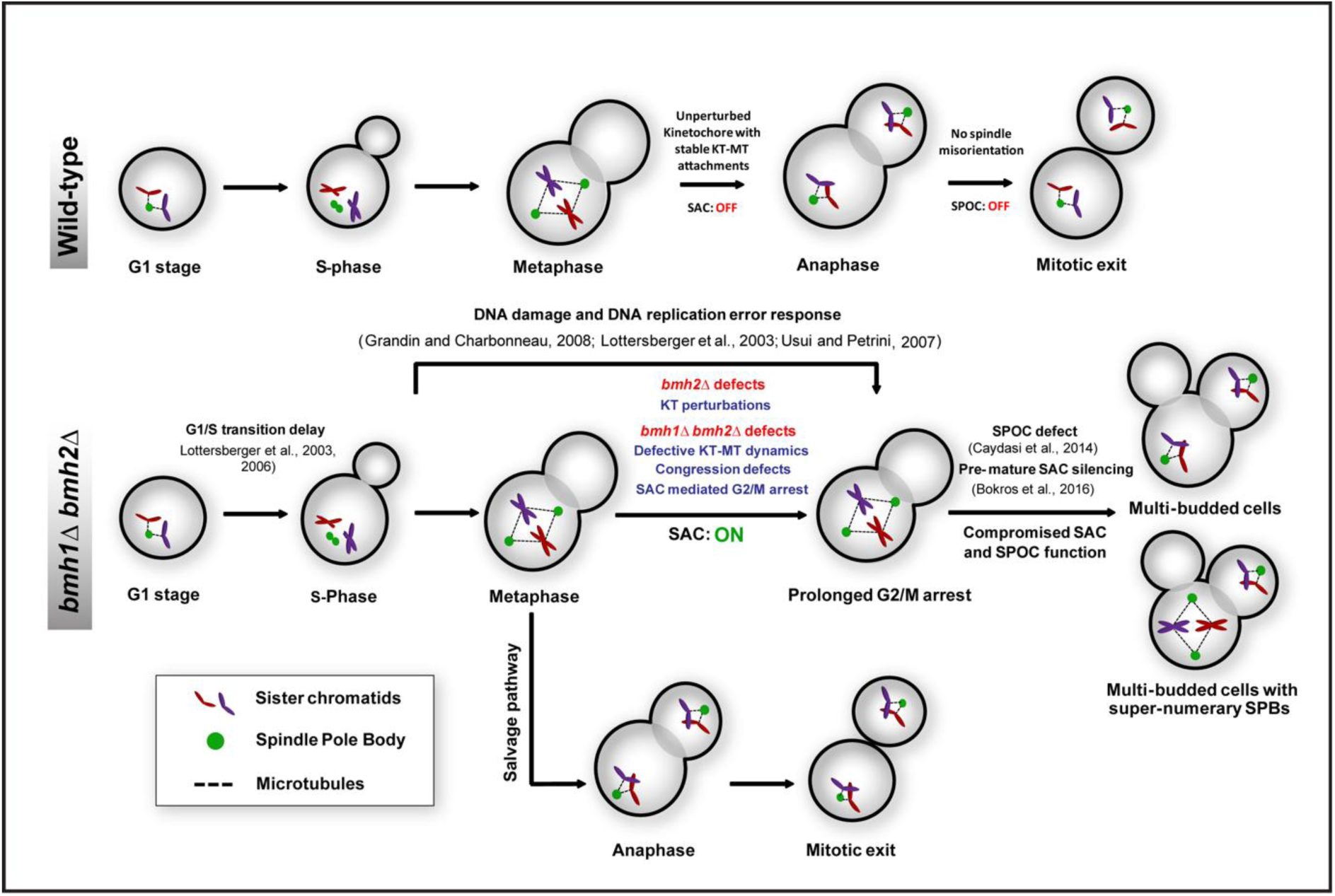
A model summarising chromosome segregation defects in *bmh* double mutants. Top, in wild-type cells, unperturbed kinetochores ensure stable KT-MT attachments and proper spindle alignment to keep SAC and SPOC off, resulting in timely G2/M transition and mitotic exit, respectively. Bottom, in the *bmh1Δ bmh2Δ* mutant cells, due to the indicated defects occurring after a transient delay in G1/S transition, SAC becomes active (SAC: ON) and arrests cells during G2/M transition. However, the presence of multi-budded cells with super-numerary SPBs denotes that a subset of cells is leaked from the arrest perhaps due to compromised SAC and SPOC functions as previously reported in *bmh1Δ* mutant. Notably, a significant fraction of *bmh1Δ bmh2Δ* mutant cells also shows no cell-cycle delay and chromosome segregation defects, indicating a possible salvage pathway exists to abrogate the defects caused by the Bmh proteins. Previously reported phenotypes (black text) of *bmh* mutants and the newly observed phenotypes (blue text) from this report are specified in the model.

To further investigate whether the Bmh proteins are targeted to the centromeres we performed chromatin immunoprecipitation (ChIP) assay. As Bmh proteins co-localize with kinetochore in all cell cycle stages (Fig 6E), asynchronously grown cultures were analyzed for centromere localization estimating centromere (*CEN)* fragment enrichment. Surprisingly, no significant localization of Bmh proteins on the centromere was observed (Fig S5A, B). Failure to detect centromeric localization by ChIP is perhaps due to the transient nature of the physical interaction between Bmh proteins and the kinetochore protein(s). In summary, we conclude that the Bmh protein either partially or completely localizes at the kinetochores across the cell-cycle stages to functionally contribute to the kinetochore functions.

## Discussion

The 14-3-3 family of proteins are conserved across eukaryotes and serve diverse regulatory functions of the cell. They exist in number of isoforms with shared and unshared functions across the eukaryotes (Kumar, 2017). Budding yeast 14-3-3 protein homolog with two isoforms (Bmh1 and Bmh2) act as regulatory molecules primarily by recognizing certain conserved phosphorylated motifs on proteins involved in various crucial activities of the cell. Among myriad other functions, Bmh proteins’ role on faithful chromosomal segregation has been reported before (Lottersberger et al., 2003; Kumar, 2018). Our further investigation in this study shows that Bmh proteins are involved in KT-MT related functions. Our findings unravel a novel function of Bmh proteins that promote kinetochore organization, stable KT-MT dynamics, and chromosome congression thus facilitating proper chromosome congression during mitosis.

### Bmh proteins contribute to the kinetochore function

The functions of the 14-3-3 proteins in the cell cycle are extensively studied across eukaryotes. Their roles in DNA replication and repair, spindle orientation, G1/S and G2/M transitions are well established (Lottersberger et al., 2003, 2006; Grandin and Charbonneau, 2008; Caydasi et al., 2014). Nevertheless, whether these proteins have roles in kinetochore function was never investigated. Previous studies on human and *Drosophilla* reported physical interaction between 14-3-3 isoforms and kinetochore proteins CENP-A and CENP-C (Goutte-Gattat et al., 2013, Przewloka et al., 2011). Additionally, in budding yeast, Bmh1 physically interacts with an intermediate filament protein Fin1, to resist its kinetochore localization which has implications in SAC silencing (Akiyoshi et al., 2009; Bokros et al., 2016, 2021). In this report, we demonstrated the functional correlation between Bmh proteins and the kinetochore function using assays including transcriptional read-through, dicentric stabilization, genetic interaction, and sub-cellular co-localization.

Since Iml3 harboring Bmh-interacting motif physically interacts with Bmh1-interacting protein Fin1 (Kakiuchi et al., 2007; Akiyoshi et al., 2009), we speculate that Bmh proteins may be targeted at or close to the centromeres through trimeric interaction of Iml3-Fin1-Bmh. Alternatively, but not mutually exclusively, Bmh proteins can be targeted at the centromeres through an association with phosphorylated H3S10 mark written by yeast aurora B kinase, Ipl1 (Jain et al., 2021). Bmh proteins prefer binding to the cruciform DNA which is present at the replication origins (Callejo et al., 2002). Although budding yeast centromeres also possess cruciform structures, direct/indirect *CEN* DNA binding of Bmh proteins is unlikely as we failed to detect them at the centromeres by ChIP. Since Bmh proteins capable of forming homo and heterodimers can cross-link two phosphoproteins, it is not surprising that Bmh proteins may have roles in proper kinetochore assembly by bridging kinetochore proteins as several of them are phosphorylated. For instance, in humans, a few 14-3-3 isoforms were reported to interact with phosphorylated CENP-A and CENP-C to form a connecting bridge and thereby stabilizes the kinetochore ensemble (Goutte-Gattat et al., 2013). Our observations of kinetochore declustering (Fig 2D) and mis-localization of the kinetochore proteins (Fig 2C) in absence of both the Bmh proteins further support this notion. Notably, while comparing the independent contributions of the Bmh isoforms, Bmh2 showed functional synergy with non-essential kinetochore proteins (Fig 5A-F). As Bmh isoforms were known to interact with phosphorylated Fin1 (Mayordomo and Sanz, 2002; Akiyoshi et al., 2009), the observed Bmh2-specific interaction with kinetochores may also be Fin1 dependent. Since early Fin1 localization and consequent SAC silencing at kinetochores were specific in *bmh1Δ* mutants (Bokros et al., 2016), it can be argued that Bmh2 perhaps holds unshared functions at kinetochores. In summary, we report here both cumulative and unshared contributions of Bmh isoforms related to kinetochore functions. In the context of unshared functions, it is interesting to observe that unlike *bmh2Δ*, *bmh1Δ* mutant alleviated the impact of non-essential kinetochore mutants (Fig 5A-F). Currently, we are uncertain about the reason. However, previously, *bmh1Δ* mutants were reported to remove inhibitory effects on the longevity factors that increase stress tolerance and extend the chronological life span of budding yeast cells (Wang et al., 2009). To correlate this phenotype of *bmh1Δ* with the effect on kinetochore requires further investigation.

### Bmh proteins function in chromosome segregation

We noticed several phenotypes of *bmh1Δ bmh2Δ* double mutant that account for the involvement of Bmh proteins in multiple events of chromosome segregation. The phenotypes include a delay in SPB-kinetochore duplication (Fig 1D), accumulation of large-budded population with well-separated SPBs and kinetochores in an unsegregated DAPI mass within mother and away from bud-neck (Fig 1C), kinetochore structural perturbations (Fig 5D, F), defective chromosome congression (Fig 3A, B) and multi-budded phenotype indicating uncoupling of chromosome segregation with the cell cycle due to checkpoint defect (Fig 4C, D).

The observed delay in SPB-kinetochore duplication, their disjunction (Fig 1D), and the accumulation of multi-budded cells (Fig 4D) in *bmh* mutants are perhaps due to the contributions of Bmh proteins in DNA replication, kinetochore functions and DNA damage checkpoint (DDC) activation, respectively (Lottersberger et al., 2003; Usui and Petrini, 2007; Grandin and Charbonneau, 2008; Bokros et al., 2016). In fission yeast, 14-3-3 homologs (Rad24 and Rad25) were shown to be necessary for DDC mediated cell cycle arrest (Ford et al., 1994; Zeng and Piwnica-Worms, 1999). In specific, Rad24 sequesters Cdc25 phosphatase from the nucleus, in response to DNA damage to arrest cells during G2/M transition. Similarly in humans, 14-3-3 isoforms have a strong role in *p53*-mediated cell cycle arrest or apoptosis in response to DNA damage (Hermeking et al., 1997; Taylor and Stark, 2001).

The increase in multi-budded cells in *bmh* double mutants upon removal of Mad2 (Fig 4D) also suggests that lack of Bmh proteins causes kinetochore perturbation resulting in altered KT-MT dynamics and SAC activation, which may also contribute to the delay in SPB-kinetochore separation (Fig 1D). In *Drosophila* oocytes, the SAC function of 14-3-3 isoforms is reported as the proteins control the kinetochore tension-sensing by controlling the localization of chromosomal passenger complex (CPC) component Borealin (Repton et al., 2022). Notably, it is possible that DNA damage generated due to improper DNA replication in *bmh1Δ bmh2Δ* double mutant might also activate the SAC since in budding yeast it has been reported that DNA damage-induced epigenetic alteration at the centromeres can alter kinetochore assembly to activate the SAC (Dotiwala et al., 2010). Overall, the phenotypes of the cell cycle delay as well as the presence of a sub-population of multi-budded cells in *bmh1Δ bmh2Δ* double mutants, suggest that the mutants not only activate DDC and/or SAC but are also compromised in sustaining the checkpoint-mediated cell cycle arrest. In support of this, it was reported earlier that *bmh* mutants are sensitive to DNA damaging and microtubule depolymerizing drugs (Grandin and Charbonneau, 2008); and *bmh1Δ* mutant, in particular, was found to cause premature SAC silencing (Bokros et al., 2016). Besides KT-MT related defects, budding yeast cells defective in kinesin motors tend to activate SAC (Jin et al., 2012; Kornakov et al., 2020; Sherwin et al., 2022). The presence of declustered kinetochores (Fig 2D) and chromatid hyperstretching phenotype (Fig 3A) in *bmh* mutants could possibly due to malfunctioning of microtubule-associated motors. Previously, in an affinity purification approach, Bmh proteins were co-purified with nuclear kinesin-related motors (Cin8, and Kar3), a Kar3 associated protein (Cik1), and protein involved in nuclear orientation and congression (Bim1) during mitosis (Kakiuchi et al., 2007). *cin8Δ* or *kar3Δ* mutant displayed kinetochore declustering phenotype similar to the *bmh* mutants (Tytell and Sorger, 2006; Jin et al., 2012) and *cik1Δ* cells, like *bmh* mutants, show activation of SAC and an increase in syntelic attachment rate with DNA replication errors (Liu et al., 2011; Jin et al., 2012). Moreover, Bim1 and Kar3 are known to have a significant role in nuclear orientation and migration during mating and mitosis (Beach et al., 2000; Molk et al., 2006). Noticeably, in a significant percentage of *bmh1Δ bmh2Δ* double mutants cells the DAPI mass (nucleus) remains away from bud-neck perhaps due to underlying motor protein-mediated nuclear migration defects (Fig 1C). Direct functional correlations between 14-3-3 isoforms and microtubule-associated proteins including motor proteins were observed in higher eukaryotes (Dorner et al., 1999; Lu and Prehoda, 2013; Beaven et al., 2017; Chen et al., 2019); however, whether this holds true in budding yeast requires further investigation. In summary, we contribute to the growing list of cell cycle functions of Bmh proteins by providing evidence that these proteins influence the kinetochore ensemble, stable KT-MT dynamics, synchronous chromosome congression and sustained activation of the checkpoints. Based on our observations and previous literature, we generated a model explaining the phenotypes of *bmh1Δ bmh2Δ* double mutants in comparison with wild-type (Fig 7).

### A possible mechanism of action of 14-3-3 proteins in kinetochore ensemble

The 14-3-3 proteins recognize their binding partners via phosphorylated motifs. Recently, these motifs were observed to be located within the intrinsically disordered regions (IDR) of the binding partners (Bustos, 2012; Sluchanko and Bustos, 2019). Amongst the 14-3-3 binding partners, numerous of them were predicted to have propensity to phase separate as membrane-less entities which might have biological significance (Huang et al., 2022) and 14-3-3 proteins may promote the propensity through binding. In this context, the literature suggests that 14-3-3 isoforms often control the functions of their binding partners by temporally sequestering them from their native cellular localization sites. For instance, 14-3-3 proteins sequester Cdc25 from the nucleus (Lopez-Girona et al., 1999; Zeng and Piwnica-Worms, 1999; Meng et al., 2013), Fin1 from kinetochore (Bokros et al., 2016), Bfa1 from SPB (Caydasi et al., 2014), Borealin and Kinesin-14/Ncd from microtubules (Repton et al., 2022) and BAD proteins from the apoptotic pathway (Tan et al., 2000). We presume that the binding partners, while under sequestration through their interaction with the Bmh proteins, may exist as phase-separated entity. In this study, we observed that some phenotypes of *bmh* mutants similar to kinetochore mutants and partial co-localization (Fig 6) between Bmh and the kinetochore proteins suggesting possible adapter function of Bmh proteins in building up kinetochore ensemble. However, we failed to detect the Bmh proteins at the centromeres by ChIP assay (Fig S5A, B). It is possible that the interaction of Bmh proteins with the centromeres is too subtle to detect with our assay. Alternatively, the function of the Bmh proteins at the kinetochore can also be manifested by sequestration of the kinetochore proteins into phase-separated entities so that the latter become available at the centromeres spatiotemporally regulating their hierarchical assembly at the kinetochore. Investigating further on Bmh proteins’ mechanism of action in regulating kinetochore integrity will be an interesting prospective. It would be intriguing to investigate in the future whether 14-3-3 proteins’ role on kinetochore is conserved across eukaryotes.

## Acknowledgement

We acknowledge the central instrumentation facility of IIT Bombay at BSBE department. SKG is supported by DST-SERB grant (CRG/2020/000444). KAG and PA are supported by DST-INSPIRE fellowship (DST/INSPIRE/03/2014/003008-IF150117) and CSIR fellowship (09/087(0972)/2019-EMR-I), respectively.

## Materials and methods

### Yeast strains and culturing methods

All *Saccharomyces cerevisiae* strains are of either SK1 or W303 genetic background. The detailed descriptions of strains and plasmids used in this study are listed in Table S1. Unless specified, as a standard condition, the strains were grown in rich medium (YEPD) at 30°C till mid-log (O.D._600_ = ∼1.0) before harvesting for the experiments. Gene modifications like C-terminal fluorescence or affinity tagging, gene deletions, and linearized plasmid integration into specific genome loci were achieved through lithium acetate transformation (Gietz and Woods, 2002) using homologous recombination. The linear cassettes for gene modifications were PCR amplified using template plasmids acquired from Euroscarf (Wach et al., 1997), and the modifications were verified by diagnostic PCR. For chromatin hyperstretching experiments, we inserted [TetO]_224_ arrays at 1.4 kb away from *CEN5* in the cells constitutively expressing TetR-GFP, as performed elsewhere (Michaelis et al., 1997; Tanaka et al., 2000). For genetic interaction with Mad2, the spores with required genotypes were obtained through meiotic induction of appropriate diploids, as described previously (Mehta et al., 2014) followed by tetrad dissection using a Zeiss Axio (Scope A1) micromanipulator (20x objective) in YEPD plates and incubation at 30°C for 2-3 days for the spore growth.

### Cell spotting and drug sensitivity assays

The cells grown in standard conditions till mid-log were sonicated briefly following which they were counted using haemocytometer, 10-fold serially diluted and spotted on control (DMSO) and benomyl-containing YEPD plates. The appropriate concentrations of benomyl (Sigma, #45339) dissolved in DMSO were added while preparing the plates. After spotting, the plates were incubated at 30°C for 2-5 days depending on the growth rate of mutants and genetic background of the strains, before they were photographed.

### Western blot and quantification

For whole-cell protein extraction, nearly 10 ml of mid-log (O.D._600_ = ∼1.0) grown cells were harvested and treated with 0.1 NaOH solution, as described before (Kushnirov, 2000). Subsequently, the processing of extracted protein samples, western blotting, and relative band intensities quantifications were performed, as mentioned elsewhere (Mittal et al., 2020). The antibodies and their dilutions (in 1X TBST buffer) were mentioned as follows. Primary antibodies: rat anti-HA (Roche, # 12158167001) at 1:5000, mouse anti-GFP (Roche, # 11814460001) at 1:3000, and rat anti-tubulin (Serotec, MCA78G) at 1:5000. Secondary antibodies: Horseradish peroxidase (HRP)-conjugated goat anti-rat (Jackson, # 112-035-167) and goat anti-mouse (Jackson, # 115-035-166) at 1:5000.

### Minichromosome stability assay

To access minichromosome (*CEN* plasmid) stability, we followed method as described earlier (Prajapati et al., 2017; Kumar et al., 2021). Typically, the cells harboring *CEN* plasmids (YCplac3, (Gietz and Sugino, 1988)) were grown overnight in selective media (SC-Ura), and re-inoculated at O.D._600_ = 0.005 into non-selective media (YEPD) and incubated at 30°C for 30-40 h to allow cells to reach ‘N’ generations. The plasmid-containing cell fractions at the initial (f0) and final (fN) time points in YEPD were estimated by plating equal volumes of cultures on SC-Ura and YEPD plates. The percentage minichromosome loss rate was calculated using the following equation, loss rate (%) = (1/N) [ln (f0/fN)] x 100; N = number of generations; f0 = fractions of plasmid-bearing cells at ‘0’ generation; fN = fractions of plasmid-bearing cells at ‘N’ generations.

### Dicentric plasmid stability

The dicentric plasmid stability assay and transcriptional read-through assay were performed, as described elsewhere (Doheny et al., 1993). To construct a dicentric plasmid pRT1 (*pGAL1-CEN4*-LacZ), a 110 bp long *CEN4* fragment was PCR amplified (using *CEN4* primers, Table S2) and cloned into SalI site placed upstream to LacZ ORF of a centromeric plasmid pAKD06 (Rizvi et al., 2017) (I mentioned Rizvi as a reference because Rizvi referred this plasmid source as ‘PJB lab BSBE IITB’) harboring *pGAL1*-LacZ cassette. As the readout for mitotic instability of dicentric plasmid was reported earlier as colony size heterogeneity on the transformant plate (Mann and Davis, 1983; Koshland et al., 1987; Doheny et al., 1993), mid-log (O.D._600_ = ∼1.0) grown cells were transformed with pRT1 plasmid, and the size heterogeneity of transformant colonies were relatively compared between wild-type and mutants on the selection plates.

### Transcriptional read-through assay

We performed transcriptional read-through assay as described elsewhere (Mann and Davis, 1983; Koshland et al., 1987; Doheny et al., 1993). We utilized the pRT1 plasmid (this study) harboring *pGAL1-CEN4*-LacZ to assess the *GAL* promoter driven LacZ expression with the transcriptional block imposed by kinetochores assembled over *CEN4*. To do so, the *pGAL1-CEN4*-LacZ cassette got excised from pRT1 plasmid and cloned into pRS406 (yeast integrative plasmid) using HindIII and SpeI restriction enzymes to construct pRT2 plasmid. The pRT2 was linearized using StuI and was integrated into *URA3* locus of wild-type and mutant cells and the transformants were maintained under galactose to keep the integrated *CEN4* (*pGAL1-CEN4*-LacZ) inactive. In the wild-type, kinetochore formed on the integrated *CEN4* offers resistance to LacZ expression which was assayed by β-galactosidase activity. In contrast, in the mutants, due to compromised kinetochore integrity, the resistance is attenuated causing increased LacZ expression. The β-galactosidase activity was determined by ONPG (o-nitrophenyl β-D-galactopyranoside) liquid assay (quantitative) as mentioned elsewhere (Clontech Laboratories, 2008). β-galactosidase activity from each sample was calculated from at least three biological replicates and the concentration of cells (O.D._600_) was normalized to compare the β-galactosidase activity between wild-type and mutants.

### Fluorescence imaging

For fluorescence imaging, as described elsewhere (Mittal et al., 2020), the cells grown in standard conditions till mid-log (O.D._600_ = ∼1.0) were fixed by adding 5% formaldehyde solution directly to the culture media and incubated at room temperature (RT) for 5 mins. The fixed cells were harvested and washed twice with 0.1 M phosphate buffer (pH 7.5) before imaging. For chromatin visualization, the cells harvested after formaldehyde fixation were briefly washed once with 50% ethanol, followed by two 0.1 M phosphate buffer (pH 7.5) washes. Finally, the cells were re-suspended in freshly prepared DAPI (Invitrogen, #D1306) (1 μg/ml) solution before imaging. Optionally, a brief sonication step was introduced to de-clump cells before imaging. The images were acquired through Zeiss Axio Observer Z1 fluorescence microscope (63× or 100x) in z-stack mode (0.2-0.5 μm spacing). For intensity comparison and 3D distance measurements, a Zeiss confocal laser scanning microscope (LSM 780) with a 32-array GaAsP detector was used. Based on the fluorescence signal intensity, the exposure time was determined and kept constant across the samples for comparison.

### Image processing

The z-stack images acquired through Zeiss Axio (Observer Z1) fluorescence microscope or confocal laser scanning (LSM 780) microscope were merged and processed for further analyses using Zeiss Zen 3.1 (blue edition) software. In represented merged images, the cell boundaries (dotted lines) were delineated referencing their brightfield images. For 3D distance measurements, the distance between any two fluorescent foci was determined by marking their centroids across Z-stacks using the Imaris 8.0.2 (Bitplane/Slice tool) software, as described elsewhere (Mittal et al., 2020). The obtained SPB-SPB and *CEN*-SPB 3D distances (in nm) were utilized to determine the cell cycle stage and *CEN*-SPB proximity, respectively; as described elsewhere (Tytell and Sorger, 2006; Sau et al., 2014).

The line scans were performed over live-cell images after merging their z-stacks projecting maximum fluorescence intensities using Zeiss Zen 3.1 (blue edition) software. The kinetochore signal distribution in the merged images was obtained using the ‘Profile’ tool in Zeiss Zen 3.1 (blue edition) software.

The Pearson’s correlation coefficient (PCC) values to assess the co-localization of any two fluorescent signals were calculated from merged z-stack images using Imaris 8.0.2 (Coloc tool) software, as described elsewhere (Prajapati et al., 2017). The obtained PCC values were categorized as mentioned earlier (Zinchuk and Grossenbacher-Zinchuk, 2014) to determine the co-localization significance.

### Time-lapse imaging

For live cell time-lapse imaging, cells were grown overnight at 30°C in YEPD media supplemented with adenine (5 mg/ml) following which they were re-inoculated into SC media and grown till O.D._600_ = 0.5. Subsequently, 0.2 – 0.5 ml of culture was added over confocal dishes (Alkali Scientific Inc) pre-coated with concanavalin A (0.25 mg/ml) solution and air-dried for 30 mins in sterile conditions for cell adherence. Unbound cells were washed away using sterile SC broth. The confocal discs were mounted over the microscope stage (temperature maintained at 30°C) to acquire automated time-lapse images at specified time intervals (at each interval, multiple fields were acquired with Z-stacks) for 12-15 h. Images were acquired using Nipkow spinning disc (Yokogawa CSU-X1 automated model) microscope equipped with an EMCCD camera. Post-acquisition processing was performed using Zeiss Zen 3.1 (blue edition) software.

### Indirect immunofluorescence

Immunofluorescence was performed, as mentioned previously (Mittal et al., 2020; Shah et al., 2023). Briefly, 5 ml of mid-log (O.D._600_ = ∼1.0) grown cells were fixed by adding 5% formaldehyde solution directly to culture media and incubated at room temperature (RT) for 15 mins. The fixed cells were harvested, washed once with PBS, resuspended in spheroplasting solution (1.2 M sorbitol, 0.1 M phosphate buffer pH 7.5), and treated with Zymolyase 20T (MP Biomedicals, #32092) (10 mg/ml) for 1 h at 30°C for spheroplasting before mounting on polylysine coated wells on a slide. After washing the unbound cells with PBS, the cells adhered to polylysine coated wells were permeabilized and flattened by immersing the slide in –20°C methanol for 5 mins and in acetone for 30 s. After allowing slides to air-dry for 1-2 mins, blocking was done using 5% skim milk solution prepared in dilution buffer (10 mg/ml BSA in PBS). Subsequently, the cells were incubated with appropriate primary and secondary antibodies and DAPI (Invitrogen, #D1306) (1 μg/ml) solutions. The intermittent washes were performed in PBS. Finally, after spreading the mounting solution (1 mg/ml phenylenediamine in 90% glycerol), the glass slides were sealed with coverslips for imaging. The antibodies and their dilutions (in dilution buffer) were mentioned as follows. Primary antibodies: rat anti-HA (Roche, # 12158167001) at 1:200, mouse anti-GFP (Roche, # 11814460001) at 1:100, mouse anti-Myc (Roche, #11667149001) at 1:200 and rat anti-tubulin (Serotec, MCA78G) at 1:5000. Secondary antibodies: (TRITC)-labeled goat anti-rat (Jackson, #115485166), Alexa Fluor 488-labeled goat anti-rat (Jackson, #112545167), and TRITC-labeled goat anti-mouse (Jackson, #115025166) at 1:200.

### Chromatin spread

Chromatin spread was performed, as mentioned previously (Prajapati et al., 2018; Mittal et al., 2020). Briefly, 2 ml of mid-log (O.D._600_ = ∼1.0) grown cells were harvested, washed once with PBS, and spheroplasted using Zymolyase 20T, as mentioned previously in the indirect immunofluorescence protocol. After spheroplasting, the reaction was stopped using an ice-cold stop solution (0.1 M Morpholineethanesulfonic acid, 1 mM EDTA, 0.5 mM MgCl_2_, 1 M sorbitol pH 6.4); and the spheroplasted cells were harvested and washed gently in PBS before mounting on a clean glass-slide. The cells were treated with freshly made paraformaldehyde solution (4% paraformaldehyde, 3.4% sucrose; add a few drops of NaOH solution to dissolve paraformaldehyde), followed by 1% Lipsol (LIP Equipment and services) solution for cell lysis. The lysed cells were homogenously spread over the slide and air-dried at RT. After overnight drying, the slides were treated with 0.4% Kodak Photoflow-200 to avoid photobleaching, followed by 5% skim milk as blocking solution. Subsequently, the slides were treated with appropriate primary and secondary antibodies and DAPI (1 μg/ml) solutions. The antibodies and their dilutions are identical to indirect immunofluorescence. The intermittent washes were performed in PBS solution. Finally, after spreading the mounting solution (1 mg/ml phenylenediamine in 90% glycerol), the glass slides were sealed with coverslips for imaging.

### ChIP assay and qPCR quantification

ChIP assay was performed, as mentioned previously (Mehta et al., 2014; Shah et al., 2023). Briefly, 50 ml of mid-log (O.D._600_ = ∼1.0) grown cells were fixed with 1% formaldehyde for 1-2 h (duration depends on chromatin proximity of the targeting protein) at 25°C. The fixed cells were harvested, washed twice with 1X TBS, and resuspended in lysis buffer (50 mM HEPES-KOH, pH 7.5, 140 mM KCl, 1 mM EDTA, 1% Triton X-100, 0.1% sodium deoxycholate, and 1X PIC (protease inhibitor cocktail, Roche)). The cells were lysed using 0.5 mm glass beads using mini-bead beater (BIOSPEC), followed by chromatin fragmentation using water bath sonicator (Diagenode SA, Picoruptor – BC100) to obtain 200-600 bps fragments. After clarifying the lysate fractions, 3-5 μg of appropriate antibodies were added and incubated at 4°C overnight with gentle rotation. Subsequently, Protein-A conjugated sepharose beads (GE Healthcare, 17-0780-01) were added and incubated at 4°C for 2 hours. Finally, the chromatin fragments were eluted by sequential washing, de-crosslinking, Proteinase K treatment, and PCI (Phenol:Chloroform:Isoamyl) purification. The enrichment of obtained chromatin fragments was estimated using qPCR (BioRad CFX96 Real-Time machine) using specific primers targeting specific and non-specific (negative control) chromatin loci, listed in Table S2. As mentioned before (Mehta et al., 2014; Mittal et al., 2020), the following equation was used to estimate the percentage chromatin enrichment. ChIP efficiency = Enrichment/Input X 100; Enrichment/Input = E˄-ΔCT; ΔCT = CT(ChIP) − [CT(Input) − LogEX(D)]; E = primer efficiency value, CT = Threshold values obtained from qPCR, D = Input dilution factor. E was estimated as {[10^(–1/slope)]– 1} from standardization graphs of CT values against dilutions of the input DNA.

### Statistical analyses

Error bars in the individual bar graphs represent the standard error (SE) derived from the standard deviation (SD) of at least three independent biological replicates. The statistical significance (*p*) was determined by two-tailed Student’s t-test (paired/un-paired, depending on the data type). The ‘N’ values denote the total number of cells analyzed, obtained from ‘n’ number of replicates of individual assay. The *p* values less than or equal to 0.05 are categorized as significant differences. The SD, SE, and statistical significance (*p*) values were calculated using automated modules of Microsoft Excel/GraphPad Prism 9.0 (Version 9.4.1) software.

## References

1. Aitken, A., Collinge, D. B., van Heusden, B. P. H., Isobe, T., Roseboom, P. H., Rosenfeld, G., et al. (1992). 14-3-3 proteins: a highly conserved, widespread family of eukaryotic proteins. Trends Biochem Sci 17. doi: 10.1016/0968-0004(92)90339-B.

2. Akiyoshi, B., Nelson, C. R., Ranish, J. A., and Biggins, S. (2009). Quantitative proteomic analysis of purified yeast kinetochores identifies a PP1 regulatory subunit. Genes Dev. doi: 10.1101/gad.1865909.

3. Beach, D. L., Thibodeaux, J., Maddox, P., Yeh, E., and Bloom, K. (2000). The role of the proteins Kar9 and Myo2 in orienting the mitotic spindle of budding yeast. Current Biology 10. doi: 10.1016/S0960-9822(00)00837-X.

4. Beaven, R., Bastos, R. N., Spanos, C., Romé, P., Fiona Cullen, C., Rappsilber, J., et al. (2017). 14-3-3 regulation of Ncd reveals a new mechanism for targeting proteins to the spindle in oocytes. Journal of Cell Biology 216. doi: 10.1083/jcb.201704120.

5. Bokros, M., Gravenmier, C., Jin, F., Richmond, D., and Wang, Y. (2016). Fin1-PP1 Helps Clear Spindle Assembly Checkpoint Protein Bub1 from Kinetochores in Anaphase. Cell Rep. doi: 10.1016/j.celrep.2016.01.007.

6. Bokros, M., Sherwin, D., Kabbaj, M. H., and Wang, Y. (2021). Yeast Fin1-PP1 dephosphorylates an Ipl1 substrate, Ndc80, to remove Bub1-Bub3 checkpoint proteins from the kinetochore during anaphase. PLoS Genet 17. doi: 10.1371/journal.pgen.1009592.

7. Braselmann, S., and McCormick, F. (1995). BCR and RAF form a complex in vivo via 14-3-3 proteins. EMBO Journal 14. doi: 10.1002/j.1460-2075.1995.tb00165.x.

8. Bustos, D. M. (2012). The role of protein disorder in the 14-3-3 interaction network. Mol Biosyst 8. doi: 10.1039/c1mb05216k.

9. Callejo, M., Alvarez, D., Price, G. B., and Zannis-Hadjopoulos, M. (2002). The 14-3-3 protein homologues from Saccharomyces cerevisiae, Bmh1p and Bmh2p, have cruciform DNA-binding activity and associate in vivo with ARS307. Journal of Biological Chemistry 277. doi: 10.1074/jbc.M202050200.

10. Caydasi, A. K., Micoogullari, Y., Kurtulmus, B., Palani, S., and Pereira, G. (2014). The 14-3-3 protein Bmh1 functions in the spindle position checkpoint by breaking Bfa1 asymmetry at yeast centrosomes. Mol Biol Cell 25, 2143–2151. doi: 10.1091/mbc.E14-04-0890.

11. Chaudhri, M., Scarabel, M., and Aitken, A. (2003). Mammalian and yeast 14-3-3 isoforms form distinct patterns of dimers in vivo. Biochem Biophys Res Commun 300. doi: 10.1016/S0006-291X(02)02902-9.

12. Cheeseman, I. M., Enquist-Newman, M., Müller-Reichert, T., Drubin, D. G., and Barnes, G. (2001). Mitotic spindle integrity and kinetochore function linked by the Duo1p/Dam1p complex. Journal of Cell Biology 152. doi: 10.1083/jcb.152.1.197.

13. Chen, Y., Chen, X., Yao, Z., Shi, Y., Xiong, J., Zhou, J., et al. (2019). 14-3-3/Tau Interaction and Tau Amyloidogenesis. Journal of Molecular Neuroscience 68. doi: 10.1007/s12031-019-01325-9.

14. Clontech Laboratories (2008). Yeast Protocols Handbook. Yeast 1.

15. Daniel, J. A., Keyes, B. E., Ng, Y. P. Y., Freeman, C. O., and Burke, D. J. (2006). Diverse functions of spindle assembly checkpoint genes in Saccharomyces cerevisiae. Genetics 172. doi: 10.1534/genetics.105.046441.

16. Doheny, K. F., Sorger, P. K., Hyman, A. A., Tugendreich, S., Spencer, F., and Hieter, P. (1993). Identification of essential components of the S. cerevisiae kinetochore. Cell 73, 761–774. doi: 10.1016/0092-8674(93)90255-O.

17. Dorner, C., Ullrich, A., Häring, H. U., and Lammers, R. (1999). The kinesin-like motor protein KIF1C occurs in intact cells as a dimer and associates with proteins of the 14-3-3 family. Journal of Biological Chemistry 274. doi: 10.1074/jbc.274.47.33654.

18. Dotiwala, F., Harrison, J. C., Jain, S., Sugawara, N., and Haber, J. E. (2010). Mad2 Prolongs DNA Damage Checkpoint Arrest Caused by a Double-Strand Break via a Centromere-Dependent Mechanism. Current Biology 20. doi: 10.1016/j.cub.2009.12.033.

19. Ford, J. C., Al-Khodairy, F., Fotou, E., Sheldrick, K. S., Griffiths, D. J. F., and Carr, A. M. (1994). 14-3-3 Protein homologs required for the DNA damage checkpoint in fission yeast. Science (1979) 265. doi: 10.1126/science.8036497.

20. Gavade, J. N., Puccia, C. M., Herod, S. G., Trinidad, J. C., Berchowitz, L. E., and Lacefield, S. (2022). Identification of 14-3-3 proteins, Polo kinase, and RNA-binding protein Pes4 as key regulators of meiotic commitment in budding yeast. Current Biology 32. doi: 10.1016/j.cub.2022.02.022.

21. Gelperin, D., Weigle, J., Nelson, K., Roseboom, P., Irie, K., Matsumoto, K., et al. (1995). 14-3-3 Proteins: Potential roles in vesicular transport and Ras signaling in Saccharomyces cerevisiae. Proc Natl Acad Sci U S A 92. doi: 10.1073/pnas.92.25.11539.

22. Ghosh, S. K., Sau, S., Lahiri, S., Lohia, A., and Sinha, P. (2004). The Iml3 protein of the budding yeast is required for the prevention of precocious sister chromatid separation in meiosis I and for sister chromatid disjunction in meiosis II. Curr Genet. doi: 10.1007/s00294-004-0516-6.

23. Gietz, R. D., and Sugino, A. (1988). New yeast-Escherichia coli shuttle vectors constructed with in vitro mutagenized yeast genes lacking six-base pair restriction sites. Gene 74, 527–534. doi: 10.1016/0378-1119(88)90185-0.

24. Gietz, R. D., and Woods, R. A. (2002). Transformation of yeast by lithium acetate/single-stranded carrier DNA/polyethylene glycol method. Methods Enzymol 350. doi: 10.1016/S0076-6879(02)50957-5.

25. Goutte-Gattat, D., Shuaib, M., Ouararhni, K., Gautier, T., Skoufias, D. A., Hamiche, A., et al. (2013). Phosphorylation of the CENP-A amino-terminus in mitotic centromeric chromatin is required for kinetochore function. Proceedings of the National Academy of Sciences. doi: 10.1073/pnas.1302955110.

26. Grandin, N., and Charbonneau, M. (2008). Budding yeast 14-3-3 proteins contribute to the robustness of the DNA damage and spindle checkpoints. Cell Cycle. doi: 10.4161/cc.7.17.6592.

27. Guarente, L. (1993). Synthetic enhancement in gene interaction: a genetic tool come of age. Trends in Genetics 9. doi: 10.1016/0168-9525(93)90042-G.

28. Hermeking, H., Lengauer, C., Polyak, K., He, T. C., Zhang, L., Thiagalingam, S., et al. (1997). 14-3-3σ is a p53-regulated inhibitor of G2/M progression. Mol Cell 1. doi: 10.1016/S1097-2765(00)80002-7.

29. Herod, S. G., Dyatel, A., Hodapp, S., Jovanovic, M., and Berchowitz, L. E. (2022). Clearance of an amyloid-like translational repressor is governed by 14-3-3 proteins. Cell Rep 39. doi: 10.1016/j.celrep.2022.110753.

30. Huang, X., Zheng, Z., Wu, Y., Gao, M., Su, Z., and Huang, Y. (2022). 14-3-3 Proteins are Potential Regulators of Liquid–Liquid Phase Separation. Cell Biochem Biophys 80. doi: 10.1007/s12013-022-01067-3.

31. Hyland, K. M., Kingsbury, J., Koshland, D., and Hieter, P. (1999). Ctf19p: A novel kinetochore protein in Saccharomyces cerevisiae and a potential link between the kinetochore and mitotic spindle. Journal of Cell Biology 145. doi: 10.1083/jcb.145.1.15.

32. Jain, N., Janning, P., and Neumann, H. (2021). 14-3-3 Protein Bmh1 triggers short-range compaction of mitotic chromosomes by recruiting sirtuin deacetylase Hst2. Journal of Biological Chemistry 296. doi: 10.1074/jbc.AC120.014758.

33. Janke, C., Ortiz, J., Lechner, J., Shevchenko, A., Shevchenko, A., Magiera, M. M., et al. (2001). The budding yeast proteins Spc24p and Spc25p interact with Ndc80p and Nuf2p at the kinetochore and are important for kinetochore clustering and checkpoint control. EMBO Journal 20. doi: 10.1093/emboj/20.4.777.

34. Jin, F., Liu, H., Li, P., Yu, H. G., and Wang, Y. (2012). Loss of function of the Cik1/Kar3 motor complex results in chromosomes with syntelic attachment that are sensed by the tension checkpoint. PLoS Genet 8. doi: 10.1371/journal.pgen.1002492.

35. Kakiuchi, K., Yamauchi, Y., Taoka, M., Iwago, M., Fujita, T., Ito, T., et al. (2007). Proteomic analysis of in vivo 14-3-3 interactions in the yeast Saccharomyces cerevisiae. Biochemistry 46, 7781–7792. doi: 10.1021/bi700501t.

36. Kornakov, N., Möllers, B., and Westermann, S. (2020). The EB1-Kinesin-14 complex is required for efficient metaphase spindle assembly and kinetochore bi-orientation. J Cell Biol 219. doi: 10.1083/jcb.202003072.

37. Koshland, D., Rutledge, L., Fitzgerald-Hayes, M., and Hartwell, L. H. (1987). A genetic analysis of dicentric minichromosomes in saccharomyces cerevisiae. Cell 48. doi: 10.1016/0092-8674(87)90077-8.

38. Kumar, D., Prajapati, H. K., Mahilkar, A., Ma, C. H., Mittal, P., Jayaram, M., et al. (2021). The selfish yeast plasmid utilizes the condensin complex and condensed chromatin for faithful partitioning. PLoS Genet 17. doi: 10.1371/journal.pgen.1009660.

39. Kumar, R. (2017). An account of fungal 14-3-3 proteins. Eur J Cell Biol 96. doi: 10.1016/j.ejcb.2017.02.006.

40. Kumar, R. (2018). Differential abundance and transcription of 14-3-3 proteins during vegetative growth and sexual reproduction in budding yeast. Sci Rep. doi: 10.1038/s41598-018-20284-6.

41. Kumar, R., Dhali, S., Srikanth, R., Ghosh, S. K., and Srivastava, S. (2014). Comparative proteomics of mitosis and meiosis in Saccharomyces cerevisiae. J Proteomics 109, 1–15. doi: 10.1016/j.jprot.2014.06.006.

42. Kushnirov, V. V. (2000). Rapid and reliable protein extraction from yeast. Yeast 16. doi: 10.1002/1097-0061(20000630)16:9<857::AID-YEA561>3.0.CO;2-B.

43. Lahiri, S., Mehta, G. D., and Ghosh, S. K. (2013). Iml3p, a component of the Ctf19 complex of the budding yeast kinetochore is required to maintain kinetochore integrity under conditions of spindle stress. FEMS Yeast Res. doi: 10.1111/1567-1364.12041.

44. Li, R., and Murray, A. W. (1991). Feedback control of mitosis in budding yeast. Cell 66. doi: 10.1016/0092-8674(81)90015-5.

45. Liu, H., Jin, F., Liang, F., Tian, X., and Wang, Y. (2011). The Cik1/Kar3 motor complex is required for the proper kinetochore-microtubule interaction after stressful DNA replication. Genetics 187. doi: 10.1534/genetics.110.125468.

46. Lopez-Girona, A., Furnari, B., Mondesert, O., and Russell, P. (1999). Nuclear localization of Cdc25 is regulated by DNA damage and a 14-3-3 protein. Nature 397. doi: 10.1038/16488.

47. Lottersberger, F., Panza, A., Lucchini, G., Piatti, S., and Longhese, M. P. (2006). The Saccharomyces cerevisiae 14-3-3 Proteins Are Required for the G1/S Transition, Actin Cytoskeleton Organization and Cell Wall Integrity. Genetics 173, 661–675. doi: 10.1534/GENETICS.106.058172.

48. Lottersberger, F., Rubert, F., Baldo, V., Lucchini, G., and Longhese, M. P. (2003). Functions of Saccharomyces cerevisiae 14-3-3 Proteins in Response to DNA Damage and to DNA Replication Stress. Genetics 165. doi: 10.1093/genetics/165.4.1717.

49. Lu, M. S., and Prehoda, K. E. (2013). A NudE/14-3-3 pathway coordinates dynein and the kinesin khc73 to position the mitotic spindle. Dev Cell 26. doi: 10.1016/j.devcel.2013.07.021.

50. Maine, G. T., Sinha, P., and Tye, B. W. (1984). Mutants of S. cerevisiae defective in the maintenance of minichromosomes. Genetics 106. doi: 10.1093/genetics/106.3.365.

51. Mann, C., and Davis, R. W. (1983). Instability of dicentric plasmids in yeast. Proc Natl Acad Sci U S A 80. doi: 10.1073/pnas.80.1.228.

52. Martens, G. J. M., Piosik, P. A., and Danen, E. H. J. (1992). Evolutionary conservation of the 14-3-3 protein. Biochem Biophys Res Commun 184. doi: 10.1016/S0006-291X(05)80046-4.

53. Masters, S. C., Pederson, K. J., Zhang, L., Barbieri, J. T., and Fu, H. (1999). Interaction of 14-3-3 with a nonphosphorylated protein ligand, exoenzyme S of Pseudomonas aeruginosa. Biochemistry 38. doi: 10.1021/bi982492m.

54. Mayordomo, I., and Sanz, P. (2002). The Saccharomyces cerevisiae 14-3-3 protein Bmh2 is required for regulation of the phosphorylation status of Fin1, a novel intermediate filament protein. Biochemical Journal 365. doi: 10.1042/BJ20020053.

55. Mehta, G. D., Agarwal, M., and Ghosh, S. K. (2014). Functional characterization of kinetochore protein, Ctf19 in meiosis I: An implication of differential impact of Ctf19 on the assembly of mitotic and meiotic kinetochores in Saccharomyces cerevisiae. Mol Microbiol 91, 1179–1199. doi: 10.1111/mmi.12527.

56. Mehta, G., Sanyal, K., Abhishek, S., Rajakumara, E., and Ghosh, S. K. (2022). Minichromosome maintenance proteins in eukaryotic chromosome segregation. BioEssays 44. doi: 10.1002/bies.202100218.

57. Meng, J., Cui, C., Liu, Y., Jin, M., Wu, D., Liu, C., et al. (2013). The Role of 14-3-3ε Interaction with Phosphorylated Cdc25B at Its Ser321 in the Release of the Mouse Oocyte from Prophase I Arrest. PLoS One 8. doi: 10.1371/journal.pone.0053633.

58. Michaelis, C., Ciosk, R., and Nasmyth, K. (1997). Cohesins: Chromosomal proteins that prevent premature separation of sister chromatids. Cell 91. doi: 10.1016/S0092-8674(01)80007-6.

59. Mittal, P., Ghule, K., Trakroo, D., Prajapati, H. K., and Ghosh, S. K. (2020). Meiosis-Specific Functions of Kinesin Motors in Cohesin Removal and Maintenance of Chromosome Integrity in Budding Yeast. Mol Cell Biol 40. doi: 10.1128/mcb.00386-19.

60. Molk, J. N., Salmon, E. D., and Bloom, K. (2006). Nuclear congression is driven by cytoplasmic microtubule plus end interactions in S. cerevisiae. Journal of Cell Biology 172. doi: 10.1083/jcb.200510032.

61. Muslin, A. J., Tanner, J. W., Allen, P. M., and Shaw, A. S. (1996). Interaction of 14-3-3 with signaling proteins is mediated by the recognition of phosphoserine. Cell 84. doi: 10.1016/S0092-8674(00)81067-3.

62. Pangilinan, F., and Spencer, F. (1996). Abnormal kinetochore structure activates the spindle assembly checkpoint in budding yeast. Mol Biol Cell 7. doi: 10.1091/mbc.7.8.1195.

63. Pearson, C. G., Yeh, E., Gardner, M., Odde, D., Salmon, E. D., and Bloom, K. (2004). Stable kinetochore-microtubule attachment constrains centromere positioning in metaphase. Current Biology 14. doi: 10.1016/j.cub.2004.09.086.

64. Poddar, A., Roy, N., and Sinha, P. (1999). MCM21 and MCM22, two novel genes of the yeast Saccharomyces cerevisiae are required for chromosome transmission. Mol Microbiol 31. doi: 10.1046/j.1365-2958.1999.01179.x.

65. Prajapati, H. K., Agarwal, M., Mittal, P., and Ghosh, S. K. (2018). Evidence of Zip1 promoting sister kinetochore mono-orientation during meiosis in budding yeast. G3: Genes, Genomes, Genetics 8. doi: 10.1534/g3.118.200469.

66. Prajapati, H. K., Rizvi, S. M. A., Rathore, I., and Ghosh, S. K. (2017). Microtubule-associated proteins, Bik1 and Bim1, are required for faithful partitioning of the endogenous 2 micron plasmids in budding yeast. Mol Microbiol. doi: 10.1111/mmi.13608.

67. Repton, C., Cullen, C. F., Costa, M. F. A., Spanos, C., Rappsilber, J., and Ohkura, H. (2022). The phospho-docking protein 14-3-3 regulates microtubule-associated proteins in oocytes including the chromosomal passenger Borealin. PLoS Genet. 18, e1009995.

68. Rizvi, S. M. A., Prajapati, H. K., Nag, P., and Ghosh, S. K. (2017). The 2-μm plasmid encoded protein Raf1 regulates both stability and copy number of the plasmid by blocking the formation of the Rep1-Rep2 repressor complex. Nucleic Acids Res 45. doi: 10.1093/nar/gkx316.

69. Roberts, R. L., Mösch, H. U., and Fink, G. R. (1997). 14-3-3 Proteins are essential for RAS/MAPK cascade signaling during pseudohyphal development in S. cerevisiae. Cell 89, 1055–1065. doi: 10.1016/S0092-8674(00)80293-7.

70. Sanyal, K., Ghosh, S. K., and Sinha, P. (1998). The MCM16 gene of the yeast Saccharomyces cerevisiae is required for chromosome segregation. Molecular and General Genetics 260. doi: 10.1007/s004380050892.

71. Sau, S., Sutradhar, S., Paul, R., and Sinha, P. (2014). Budding yeast kinetochore proteins, Chl4 and Ctf19, are required to maintain SPB-centromere proximity during G1 and late anaphase. PLoS One 9. doi: 10.1371/journal.pone.0101294.

72. Shah, S., Mittal, P., Kumar, D., Mittal, A., and Ghosh, S. K. (2023). Evidence of kinesin motors involved in stable kinetochore assembly during early meiosis. Mol Biol Cell 34. doi: 10.1091/mbc.E22-12-0569.

73. Sherwin, D., Huetteman, A., and Wang, Y. (2022). Yeast kinesin-5 motor protein CIN8 promotes accurate chromosome segregation. Cells 11, 2144.

74. Sikorski, R. S., and Hieter, P. (1989). A system of shuttle vectors and yeast host strains designed for efficient manipulation of DNA in Saccharomyces cerevisiae. Genetics 122. doi: 10.1093/genetics/122.1.19.

75. Slubowski, C. J., Paulissen, S. M., and Huang, L. S. (2014). The GCKIII kinase Sps1 and the 14-3-3 Isoforms, Bmh1 and Bmh2, cooperate to ensure proper sporulation in Saccharomyces cerevisiae. PLoS One 9. doi: 10.1371/journal.pone.0113528.

76. Sluchanko, N. N., and Bustos, D. M. (2019). “Intrinsic disorder associated with 14-3-3 proteins and their partners,” in Progress in Molecular Biology and Translational Science doi: 10.1016/bs.pmbts.2019.03.007.

77. Sluchanko, N. N., and Gusev, N. B. (2017). Moonlighting chaperone-like activity of the universal regulatory 14-3-3 proteins. FEBS Journal 284. doi: 10.1111/febs.13986.

78. Stanhill, A., Schick, N., and Engelberg, D. (1999). The Yeast Ras/Cyclic AMP Pathway Induces Invasive Growth by Suppressing the Cellular Stress Response. Mol Cell Biol 19. doi: 10.1128/mcb.19.11.7529.

79. Tan, Y., Demeter, M. R., Ruan, H., and Comb, M. J. (2000). BAD Ser-155 phosphorylation regulates BAD/Bcl-XL interaction and cell survival. Journal of Biological Chemistry 275. doi: 10.1074/jbc.M004199200.

80. Tanaka, T., Fuchs, J., Loidl, J., and Nasmyth, K. (2000). Cohesin ensures bipolar attachment of microtubules to sister centromeres and resists their precocious separation. Nat Cell Biol 2. doi: 10.1038/35019529.

81. Taylor, W. R., and Stark, G. R. (2001). Regulation of the G2/M transition by p53. Oncogene 20. doi: 10.1038/sj.onc.1204252.

82. Thrower, D. A., and Bloom, K. (2001). Dicentric chromosome stretching during anaphase reveals roles of Sir2/Ku in chromatin compaction in budding yeast. Mol Biol Cell 12. doi: 10.1091/mbc.12.9.2800.

83. Tytell, J. D., and Sorger, P. K. (2006). Analysis of kinesin motor function at budding yeast kinetochores. Journal of Cell Biology 172. doi: 10.1083/jcb.200509101.

84. Usui, T., and Petrini, J. H. J. (2007). The Saccharomyces cerevisiae 14-3-3 proteins Bmh1 and Bmh2 directly influence the DNA damage-dependent functions of Rad53. Proc Natl Acad Sci U S A 104. doi: 10.1073/pnas.0611259104.

85. Van Heusden, G. P. H., Griffiths, D. J. F., Ford, J. C., Chin-A-Woeng, T. F. C., Schrader, P. A. T., Carr, A. M., et al. (1995). The 14-3-3 Proteins Encoded by the BMH1 and BMH2 Genes are Essential in the Yeast Saccharomyces cerevisiae and Can be Replaced by a Plant Homologue. Eur J Biochem 229. doi: 10.1111/j.1432-1033.1995.0045l.x.

86. van Heusden, G. P. H., Wenzel, T. J., Lagendijk, E. L., de Steensma, H. Y., and van den Berg, J. A. (1992). Characterization of the yeast BMH1 gene encoding a putative protein homologous to mammalian protein kinase II activators and protein kinase C inhibitors. FEBS Lett 302. doi: 10.1016/0014-5793(92)80426-H.

87. Wach, A., Brachat, A., Alberti-Segui, C., Rebischung, C., and Philippsen, P. (1997). Heterologous HIS3 marker and GFP reporter modules for PCR-targeting in Saccharomyces cerevisiae. Yeast 13. doi: 10.1002/(SICI)1097-0061(19970915)13:11<1065::AID-YEA159>3.0.CO;2-K.

88. Wang, C., Skinner, C., Easlon, E., and Lin, S. J. (2009). Deleting the 14-3-3 protein Bmh1 extends life span in Saccharomyces cerevisiae by increasing stress response. Genetics 183. doi: 10.1534/genetics.109.107797.

89. Wang, Y., and Burke, D. J. (1995). Checkpoint genes required to delay cell division in response to nocodazole respond to impaired kinetochore function in the yeast Saccharomyces cerevisiae. Mol Cell Biol 15. doi: 10.1128/mcb.15.12.6838.

90. Wargacki, M. M., Tay, J. C., Muller, E. G., Asbury, C. L., and Davis, T. N. (2010). Kip3, the yeast kinesin-8, is required for clustering of kinetochores at metaphase. Cell Cycle 9. doi: 10.4161/cc.9.13.12076.

91. Wilkert, E. W., Grant, R. A., Artim, S. C., and Yaffe, M. B. (2005). A Structural Basis for 14-3-3σ Functional Specificity,. Journal of Biological Chemistry 280, 18891–18898. doi: 10.1074/JBC.M500982200.

92. Xiangwei, H., Saurabh, A., and Sorger, P. K. (2000). Transient sister chromatid separation and elastic deformation of chromosomes during mitosis in budding yeast. Cell 101. doi: 10.1016/s0092-8674(00)80888-0.

93. Yaffe, M. B., Rittinger, K., Volinia, S., Caron, P. R., Aitken, A., Leffers, H., et al. (1997). The structural basis for 14-3-3:phosphopeptide binding specificity. Cell 91. doi: 10.1016/S0092-8674(00)80487-0.

94. Yahyaoui, W., Callejo, M., Price, G. B., and Zannis-Hadjopoulos, M. (2007). Deletion of the cruciform binding domain in CBP/14-3-3 displays reduced origin binding and initiation of DNA replication in budding yeast. BMC Mol Biol 8. doi: 10.1186/1471-2199-8-27.

95. Yahyaoui, W., and Zannis-Hadjopoulos, M. (2009). 14-3-3 proteins function in the initiation and elongation steps of DNA replication in Saccharomyces cerevisiae. J Cell Sci 122. doi: 10.1242/jcs.044677.

96. Zeng, Y., and Piwnica-Worms, H. (1999). DNA Damage and Replication Checkpoints in Fission Yeast Require Nuclear Exclusion of the Cdc25 Phosphatase via 14-3-3 Binding. Mol Cell Biol 19. doi: 10.1128/mcb.19.11.7410.

97. Zinchuk, V., and Grossenbacher-Zinchuk, O. (2014). Quantitative colocalization analysis of fluorescence microscopy images. Curr Protoc Cell Biol 62, 4.19.1–4.19.14.

