## Supplementary files for "Evidence of 14-3-3 proteins contributing to kinetochore integrity and chromosome congression during mitosis"

**<u>Supplementary data:</u>**

**Table S1: List of strains and plasmids used in this study:**

All the SK1 strains used were derived from the haploid strains with the following genotypes.

*MATa, ho::Lys2, lys2, ura3, leu2::hisG, his3::hisG, trp1::hisG*

*MATα, ho::Lys2, lys2, ura3, leu2::hisG, his3::hisG, trp1::hisG*

| **Name** | **Genotype** | **Background** | **Source** |
|---|---|---|---|
| SGY167 | *MATa, ctf19Δ::KanMx* | SK1 | This study |
| SGY824 | *MATα, leu2::tetR-GFP::LEU2,CENV::tetO::HIS3, bmh1Δ::KanMx* | SK1 | This study |
| SGY822 | *MATα, leu2::tetR-GFP::LEU2, CENV::tetO::HIS3, bmh2Δ::URA3* | SK1 | This study |
| SGY833 | *MATa, bmh2Δ::HIS3, bmh1Δ::KanMx* | SK1 | (Kumar, 2018) |
| SGY7134 | *MATa, iml3Δ::KanMx* | SK1 | This study |
| SGY1337 | *MATa, NDC80-CFP::HIS3, SPC42-EGFP::TRP1* | SK1 | This study |
| SGY834 | *MATa, bmh2Δ::URA3, NDC80-CFP::HIS3, SPC42-EGFP::TRP1* | SK1 | (Kumar, 2018) |
| SGY7156b | *MATa, bmh1Δ::KanMx, bmh2Δ::URA3, NDC80-CFP::HIS3 SPC42-EGFP::TRP1* | SK1 | This study |
| SGY 7158 | *MATa, MTW1-EGFP::TRP1 SPC42-mCherry::KanMx* | SK1 | This study |
| SGY 7159 | *MATa, bmh1Δ::KanMx, MTW1-EGFP::TRP1, SPC42-mCherry::KanMx* | SK1 | This study |
| SGY 7160 | *MATa, bmh2Δ::HIS3, MTW1-EGFP::TRP1 SPC42-mCherry::KanMx* | SK1 | This study |
| SGY 7161 | *MATa, bmh1Δ::KanMx, bmh2Δ::HIS3, MTW1-EGFP::TRP1 SPC42-mCherry::KanMx* | SK1 | This study |

| | | | |
|---|---|---|---|
| SGY 7167 | *MATa, leu2::tetR-GFP::LEU2, CENV::tetO::HIS3, SPC42-mCherry::KanMx* | SK1 | This study |
| SGY 7168 | *MATa, bmh1Δ::KanMx, leu2::tetR-GFP::LEU2, CENV::tetO::HIS3, SPC42-mCherry::KanMx* | SK1 | This study |
| SGY 7169 | *MATa, bmh2Δ::URA3, leu2::tetR-GFP::LEU2, CENV::tetO::HIS3, SPC42-mCherry::KanMx* | SK1 | This study |
| SGY 7170 | *MATa, bmh1Δ::KanMx, bmh2Δ::URA3, leu2::tetR-GFP::LEU2, CENV::tetO::HIS3, SPC42-mCherry::KanMx* | SK1 | This study |
| SGY 7162 | *MATa, mad2Δ::KanMx* | SK1 | This study |
| SGY 7163 | *MATa, bmh1Δ::KanMx, mad2Δ::KanMx* | SK1 | This study |
| SGY 7164 | *MATa, bmh2Δ::URA3, mad2Δ::KanMx* | SK1 | This study |
| SGY 7165 | *MATa, bmh1Δ::KanMx, bmh2Δ::URA3, mad2Δ::KanMx* | SK1 | This study |
| SGY 7135 | *MATa, iml3Δ::KanMx, bmh1Δ::KanMx* | SK1 | This study |
| SGY 7136 | *MATa, iml3Δ::KanMx, bmh2Δ::URA3* | SK1 | This study |
| SGY 7138 | *MATa, iml3Δ::KanMx bmh1Δ::KanMx, bmh2Δ::URA3* | SK1 | This study |
| SGY116 | *MATa/α, leu2::tetR-GFP::LEU2/leu2::tetR-GFP::LEU2 TetO-HIS3/-* | SK1 | (Mehta et al., 2014) |
| CRY1 | *MATa, leu2-3,112 trp1-1 can1-100 ura3-1 ade2-1 his3-11,15* | W303 | Yeast Resource center (YRC) |
| SGY10100 | *MATa, NDC80-6HA::KanMx, BMH1-9MYC::TRP1* | W303 | This study |
| SGY10103 | *MATa, NDC80-6HA::KanMx, BMH2-13MYC::HIS3* | W303 | This study |
| SGY10106 | *MATa, NDC80-6HA::KanMx* | W303 | This study |
| SGY7010 | *MATa, ctf19Δ::KanMx* | W303 | This study |

| | | | |
|---|---|---|---|
| SGY7011 | *MATa, bmh1Δ::LEU2* | W303 | This study |
| SGY7013 | *MATa, bmh2Δ::HIS3* | W303 | This study |
| SGY7149 | *MATa, ctf19Δ::KanMx4 bmh1Δ::LEU2* | W303 | This study |
| SGY7150 | *MATa, ctf19Δ::KanMx4 bmh2Δ::HIS3* | W303 | This study |
| SGY7181 | *MATa, leu2-3,112 trp1-1 can1-100 ura3-1 ade2-1 his3-11,15 / ura3::pRT2::URA3* | W303 | This study |
| SGY7182 | *MATa, ctf19Δ::KanMx4 / ura3::pRT2::URA3* | W303 | This study |
| SGY7183 | *MATa, bmh1Δ::LEU2 / ura3::pRT2::URA3* | W303 | This study |
| SGY7184 | *MATa, bmh2Δ::HIS3 / ura3::pRT2::URA3* | W303 | This study |

**Plasmids and their description:**

| **Name** | **Description** | **Source** |
|---|---|---|
| yCPlac33 | Yeast centromeric plasmid used for minichromosome stability assay | (Gietz and Sugino, 1988) |
| pRS406 | Yeast integrative plasmid used to integrate '*pGAL1-CEN4*-LacZ' at *URA3* locus | (Sikorski and Hieter, 1989) |
| pAKD06 | Centromeric plasmid containing '*pGAL1*-LacZ' construct. | (Rizvi et al., 2017) |
| pRT1 | Plasmid used for dicentric stability and transcriptional read-through assay | This study |
| pRT2 | Modified pRS406 containing '*pGAL1-CEN4*-LacZ' construct | This study |

1 **Table S2: List of primers used in ChIP-qPCR:**

| **Name** | **Description** | **Primer sequence (5′-3′)** |
|---|---|---|
| *CEN3* | Forward | gatcagcgccaaacaatatgg |

| | | |
|---|---|---|
| | Reverse | aacttccaccagtaaacgttt |
| *CEN4* | Forward | gcttgcaaaaggtcacatgc |
| | Reverse | gagcaggttttatgtttcgg |
| *TUB2* | Forward | cttgtagacagcgtcatgg |
| | Reverse | cagatgtcataaagtgcttcg |

### Supplementary figures:

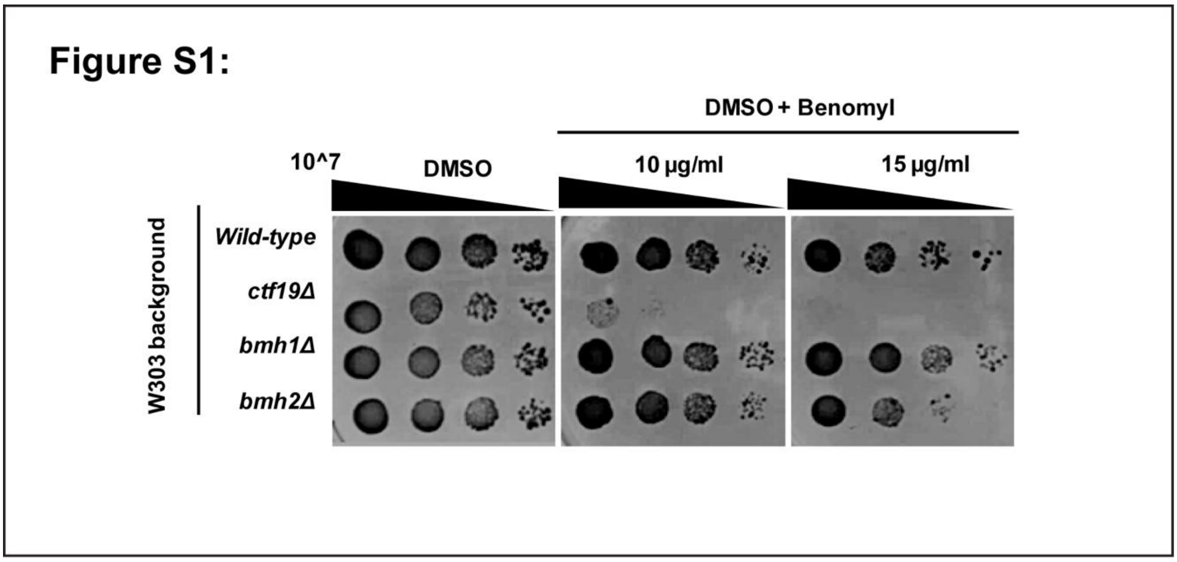


**Figure S1: Hyper-sensitivity of the *bmh* mutants to the microtubule-disrupting drug.** Wild-type and *bmh* single (*bmh1*Δ, *bmh2*Δ) mutant cells with W303 genetic background (in which *bmh* double mutant is inviable) were spotted on control (DMSO) and indicated concentrations of drug (DMSO+Benomyl) containing YEPD plates. The non-essential kinetochore mutant (*ctf19*Δ) was taken as a positive control for benomyl hyper-sensitivity. Approximately 10^7 cells were serially diluted 10-fold and spotted on the above-mentioned plates and were incubated at 30°C for 24-48 hours before imaging. Experimental replicates, n = 3.

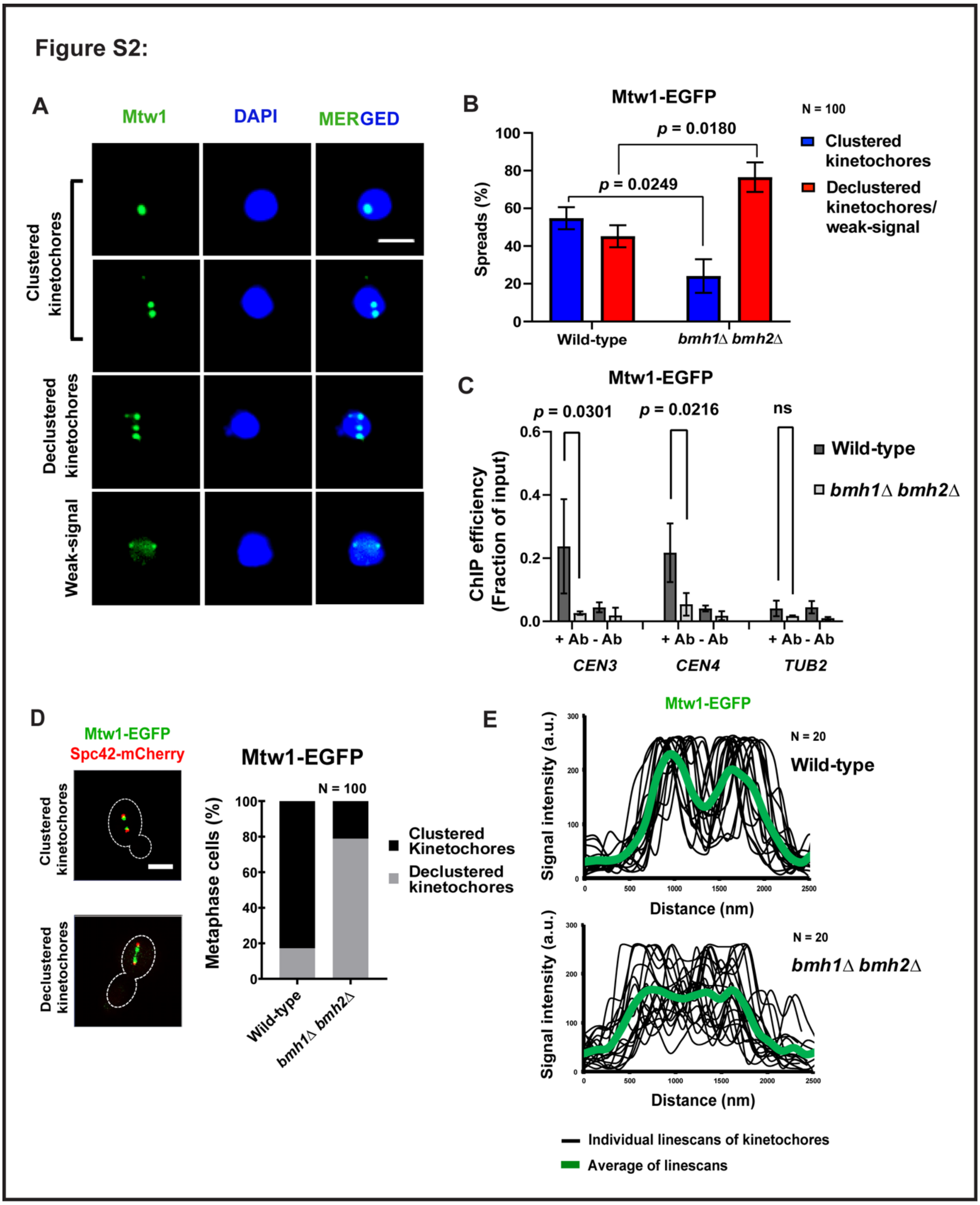


**Figure S2: Perturbed clustering and integrity of the kinetochores (Mtw1) in the absence of Bmh proteins. (A)** The representative images of the chromatin spreads to visualize kinetochore protein Mtw1 fused to EGFP in wild-type and *bmh1*Δ *bmh2*Δ cells before anaphase (undivided DAPI mass). Anti-GFP antibodies and DAPI staining were used to probe Mtw1-EGFP and chromatin, respectively. Scale bar = 2 μm. **(B)** The percentage of different categories of the spreads as mentioned in A, for the indicated strains. N = 100, Experimental replicates, n = 2. **(C)** ChIP-qPCR analyses were performed to quantify the association of Mtw1-EGFP with *CEN3, CEN4,* and *TUB2* (negative control) loci in the indicated strains. Anti-GFP antibodies were used for pull-down from the asynchronously grown mid-log cells. Error bars indicate standard error. Experimental replicates, n = 3. Statistical significance (*p*) was calculated using the two-tailed student's t-test. 'ns' represents statistically not significant. **(D)** Left, representative live cell images showing localization pattern of Mtw1-EGFP during metaphase (SPB-SPB distance = 1.5 – 2.5 μm). Scale bar = 2 μm. Right, the percentage of metaphase cells depicting 'clustered' or 'declustered' kinetochores were shown for the indicated strains. N = 100, Experimental replicates, n = 2. (E) The intensity graphs depict the distribution of fluorescence signal intensity of Mtw1-EGFP (Green) in the indicated strains. Line scans were performed along the axis between two SPBs for 20 cells to estimate mean signal intensity distribution (arbitrary units) for each strain. Experimental replicates, n = 2.

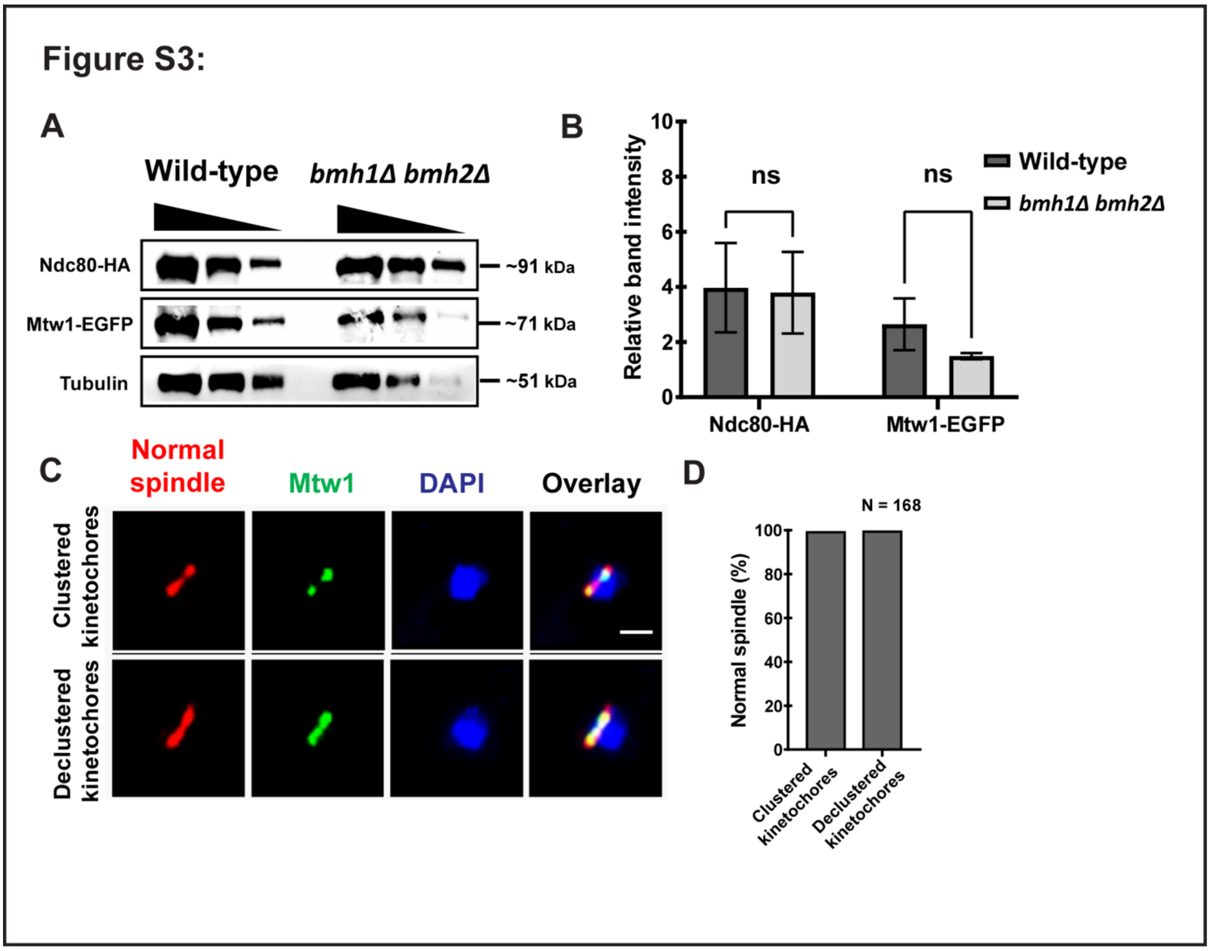


**Figure S3: The loss of Bmh proteins does not affect expression of the kinetochore proteins (Ndc80, Mtw1) and spindle morphology. (A)** Western-blots show the levels (in serial dilutions) of the indicated kinetochore proteins in asynchronously grown wild-type and *bmh1*Δ *bmh2*Δ cells. Anti-HA, anti-GFP, and anti-tubulin antibodies were used to detect the corresponding proteins. **(B)** The graph shows the relative band intensity measured from the average of first two bands intensities of the indicated proteins normalized with the background and tubulin band. Error bars indicate standard error. Experimental replicates, n = 2. Statistical significance (*p*) was calculated using the two-tailed student's t-test. 'ns' represents statistically not significant. **(C)** Indirect immunofluorescence assay to visualize metaphase spindle in cells containing clustered and declustered kinetochores. Anti-tubulin, anti-GFP, and DAPI staining were used to probe microtubule spindle, kinetochore (Mtw1-EGFP), and chromatin, respectively, in the strains indicated in (A). Scale bar = 2 μm. **(E)** The percentage of metaphase cells containing normal spindle, as shown in (C), with either clustered or declustered kinetochores. N = 160, Experimental replicates, n = 2.

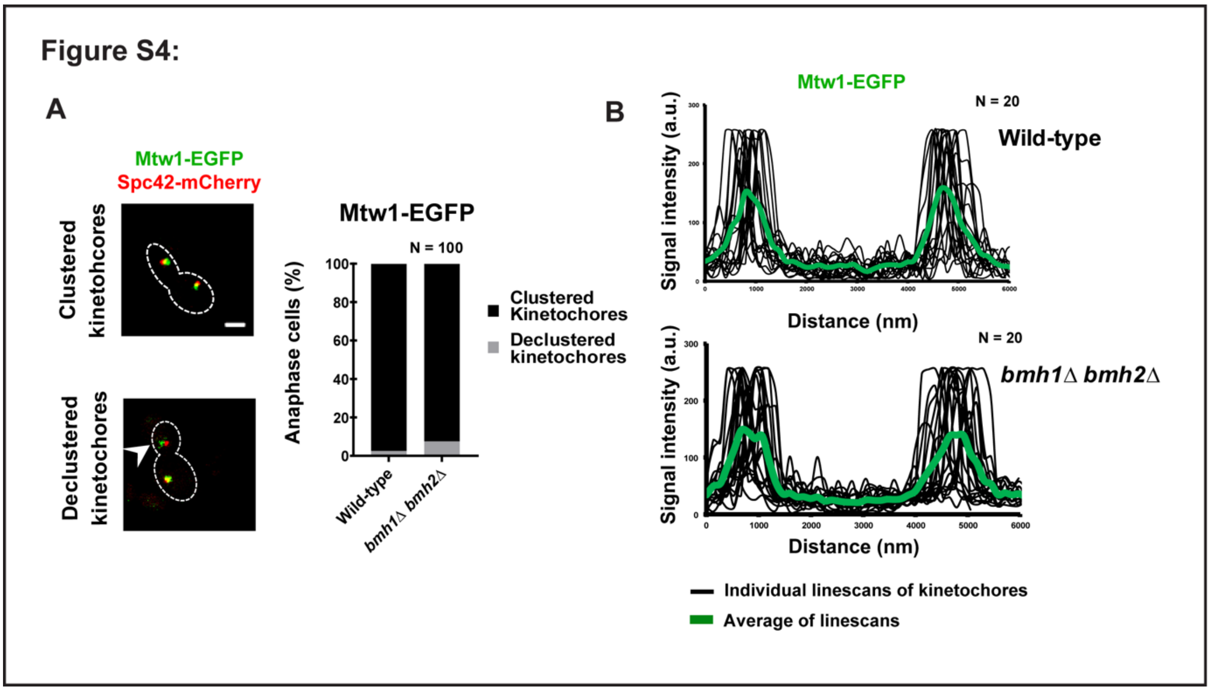


**Figure S4: kinetochores remain properly clustered during post-metaphase in *bmh* double mutant. (A)** Left, representative live cell images showing localization pattern of Mtw1-EGFP during anaphase (SPB-SPB distance > 4.0 μm). Scale bar = 2 μm. Right, the percentage of anaphase cells depicting 'clustered' or 'declustered' kinetochores were shown for the indicated strains. Noticeably, the declustering (arrowhead) during anaphase is not as prominent in metaphase cells. N = 100, Experimental replicates, n = 2. **(B)** The intensity graphs depict the distribution of fluorescence signal intensity of Mtw1-EGFP (Green) in the indicated strains. Line scans were performed along the axis between two SPBs for 20 cells to estimate mean signal intensity distribution (arbitrary units) for each strain. Experimental replicates, n = 2.

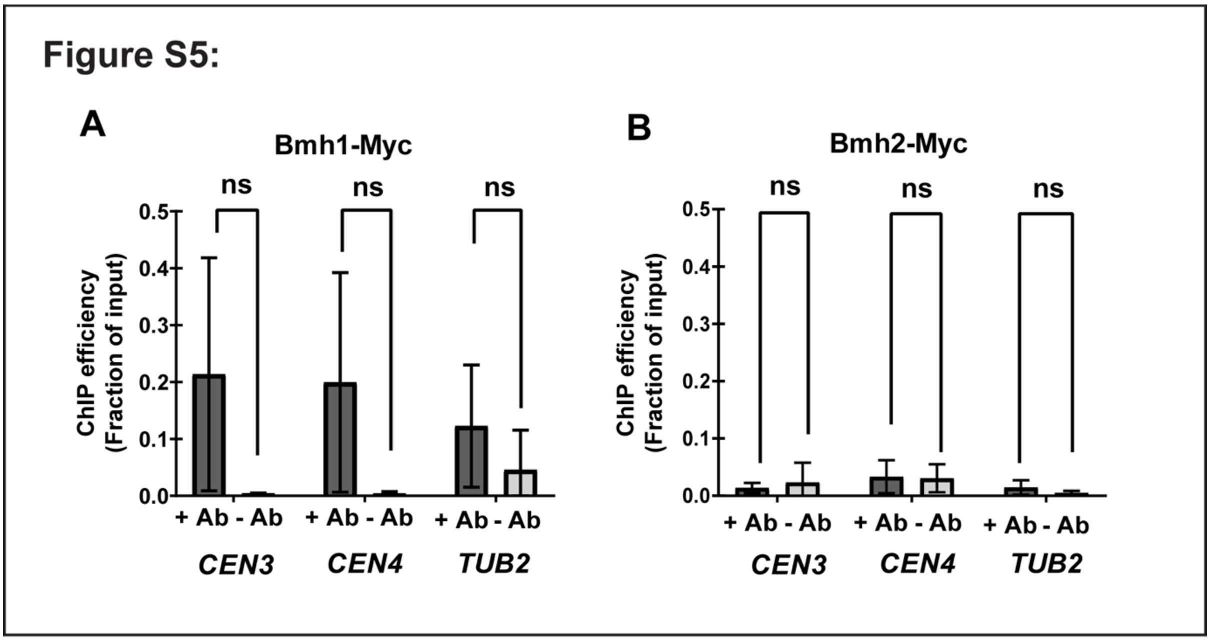


**Figure S5: Localization of Bmh proteins at the centromeres.** ChIP-qPCR analyses were performed to quantify the association of Bmh proteins (Bmh1/Bmh2) with centromere loci in the indicated strains. Anti-Myc antibodies were used to pull-down individual Bmh protein (Bmh1-Myc/Bmh2-Myc) from the asynchronously grown mid-log cells to quantify percentage enrichments of *CEN3, CEN4,* and *TUB2* (negative control) fragments. Error bars indicate standard error. Experimental replicates, n = 3. Statistical significance (*p*) was calculated using the two-tailed student's t-test. 'ns' represents statistically not significant.
